# Genomic stress drives activation of interferon signaling and innate immune pathways during SMA Type II myoblasts differentiation

**DOI:** 10.64898/2025.12.23.696216

**Authors:** Sabrina Mazzucchi, Anne Bigot, Fiorella Carla Grandi, Piera Smeriglio

## Abstract

Spinal muscular atrophy (SMA) is a neuromuscular disorder caused by loss of the *SMN1* gene. The SMN protein is ubiquitously expressed and has several roles in the cell, including the regulation of RNA metabolism and genome stability. Although SMA is primarily considered a motor neuron disease, because of its ubiquitous expression, loss of SMN leads to systemic pathological consequences, including in skeletal muscle. To better understand the cell-intrinsic effects of SMN loss on myoblast differentiation, we characterized three human SMA Type II myoblast cell lines and controls *in vitro*. We observed impaired myogenic differentiation, including reduced fusion index and nuclei alignment. RNA sequencing analysis on the SMA myoblasts revealed activation of DNA damage pathways and innate immune activation, confirmed by the presence of 53BP1 foci and increased phosphorylation of the H2AX histones, as well as accumulation of R-loop structures and an increase of single-stranded DNA in the cytoplasm. Resolution of R-loops and DNA damage is known to cause the presence of immunogenic DNA: RNA species in the cytoplasm, activating sterile inflammation pathways. Treating control myoblasts with double-stranded DNA or a cGAS agonist phenocopied the differentiation defects in SMA myotubes.

Conversely, treating SMA myoblasts with a STING inhibitor improved fusion. In summary, our results demonstrate that SMN loss exerts a cell-intrinsic impairment in muscle precursor cells, suggesting that the observed muscle dysfunction in Type II patients is not only a consequence of impaired innervation but also of the sterile inflammation caused by the genomic instability after SMN loss.

## INTRODUCTION

Spinal Muscular Atrophy (SMA) is an autosomal, recessive, developmental disorder characterized by the selective degeneration and death of motor neurons and progressive muscle atrophy [1]. SMA is caused by the homozygous loss of the Survival Motor Neuron gene (*SMN1*), which is ubiquitously expressed in the body and plays a wide variety of roles in the cell. SMA was initially solely considered a neurodegenerative disease; however, recent studies have suggested that several organ systems are affected by and involved in the disease [2,3], and may cause its complex and heterogeneous presentation. Indeed, these findings are in line with the ubiquitous nature of the SMN protein and its multiplicity of roles [4].

Despite decades of research and the existence of SMN-based therapies [5], the exact ways that SMN loss affects each cell type and tissue remain poorly understood. The *SMN1* gene codes for a 37 KDa protein, which is implicated in many different cellular processes, the most well-studied of these being its presence in snRNPs and implication in U12 splicing [6]. The complete absence of the SMN protein is lethal during embryogenesis in mice [7]. However, the human genome contains a second SMN copy, *SMN2*, which contains a single base pair modification that provides an alternatively spliced mRNA isoform lacking exon 7 (*SMN*Δ*7*). Therefore, while the *SMN1* gene can produce a full-length functional protein, nearly 90% of *SMN2* transcripts are truncated and generate a non-functional protein, termed SMNΔ7 [1]. The 10% full-length transcripts can compensate for some but not all the functions of *SMN1* and thus patients lacking *SMN1* will eventually manifest symptoms of SMA, and the severity of these symptoms will be determined by the number of *SMN2* copies [1].

Among the many tissues affected in SMA patients are skeletal muscles [8]. Initially, skeletal muscle defects were considered merely a consequence of denervation due to the death of the motor neurons [9]. However, recent studies suggest that SMN loss may also result in cell-intrinsic defects in skeletal muscle. Our previous study of paravertebral muscle from Type II patients treated with Nusinersen and Risdiplam, two drugs designed to increase the levels of the correctly spliced *SMN2* gene, showed that patients had larger myofibers with centralized nuclei, loss of oxidative phosphorylation machinery, decreased mitochondrial DNA copy number, and increased fibrosis [10]. Similar findings have been seen in muscles of other SMA patient populations, including defects observed in myoblast fusion in foetal SMA Type I samples [11]. Collectively, this has led to the hypothesis that the cell-intrinsic roles of SMN in muscle cells may also contribute to SMA muscle pathology, beyond its roles in motor neurons and the associated denervation. This has been confirmed by several muscle-cell specific knockout (KO) models, including *PAX7*, *HSA, and MYOD1* models, which show muscle-specific disease [8,12–14].

In addition, these *in vivo* studies have been supported by various *in vitro* studies with mouse and human muscle cells. *In vitro* studies represent a genetically tractable system to study the cell-instinct roles of SMN on early human myogenesis. Co-cultures of muscle satellite cells (MuSCs) derived from SMA patients with control motor neurons showed impaired innervation, suggesting an SMN muscle-specific role in the formation of the neuromuscular junctions [15]. Myoblasts from Type I patients form smaller and disorganized myotubes, with delayed maturation and decreased acetylcholine receptor expression [16] and have metabolic dysfunction [17]. This has also been recapitulated in C2C12 cells after SMN knockdown [18,19]. However, although this *in vivo* and *in vitro* work in muscle has been seminal in highlighting the importance of SMN in muscle tissue and muscle cells, including satellite cells and myoblasts, these studies did not fully resolve what cellular roles SMN plays in muscle cells, and how this might differ from its roles in motor neurons.

The primary function of SMN remains a complex and unresolved question. SMN has been linked to several processes in the cell, including cytoskeletal and actin remodelling, calcium regulation, splicing, ribosome assembly, and telomerase biogenesis, among others [4]. More recently, SMN has been proposed to be involved in several chromatin and transcription regulating processes, using the methylation binding functions of its TUDOR domain [20,21]. For example, SMN can bind methylated lysine residues on histone H3 [21], as well as the symmetric demethylation of arginine on the carboxyl-terminal domain of RNA Poll II subunit *POLR2A* [22,23]. The latter is thought to mediate the SMN-dependent calling of SETX and ZPR1 to the RNA Pol II complex [24,25]. The absence of SMN in this scenario causes the accumulation of co-transcriptional R-loops [24], DNA:RNA structures formed during transcription. In the case of motor neurons, the accumulation of these R-loops is thought to cause DNA damage and cell death [26,27], although the directionality of this relationship remains unresolved. Indeed, SMN loss is associated with genome instability and increased DNA damage in several tissues, and this signature can also be seen in patient muscle samples [10].

Distinguishing intrinsic muscle alterations from the effects of denervation and impaired motor neuron signalling is often challenging; however, such work is critical, as understanding the alterations that occur in SMA muscle is an important clinical priority to address the remaining peripheral tissue defects in SMA, even with the presence of three disease-modifying therapies. Thus, further studies are needed to elucidate the cell-intrinsic defects caused by SMN deficiency that underlie the muscle-specific pathology in SMA. With this goal in mind, in this work, we sought to use patient-derived immortalized Type II SMA myoblasts to study the effect of *SMN1* loss on Type II SMA myoblast differentiation. We find that the increased genomic instability of SMA myoblasts, including the accumulation of both R-loops and DNA damage, can trigger innate immune signalling and interferon signalling, which causes defective and disorganized myotube formation.

## RESULTS

### Immortalized SMA II myoblasts show reduced and disorganized myotube formation

To analyze the cell-intrinsic consequences of low SMN levels on myotube formation, independently of the motor neuron death, we used an *in vitro* model of muscle progenitor cells, myoblasts, immortalized from the surgical discard of Type II SMA patients and controls without neuromuscular disease, as previously described [28]. We assembled a cohort of three SMA lines (SMAII.1, SMAII.2, and SMAII.3), whose average donor age was 10 years old and three control cell lines from muscle without neuromuscular disease (C.1, C.2, C.3), with an average donor age of 22 (**Table 1**). We validated the *SMN1* and *SMN2* copy number by qPCR and observed, as expected, no signal from the *SMN1* gene, and an increased signal from *SMN2* (**Supplemental Fig.1 A-B**). To verify the total *SMN1*+*SMN2* copies, we performed digital droplet PCR (ddPCR) and observed that SMAII.1, SMAII.2, and SMAII.3 had three *SMN* copies on average, and controls had four (**Supplemental Fig.1 C**). This corresponds to the average *SMN2* copy number observed in SMA type II patients [1]. SMN western blot analysis showed a 3-fold decrease in SMN protein expression in all SMA cell lines (**Supplemental Fig.1 D-E**).

**Table 1:** Biopsy information for immortalized myoblast cell lines used in this study.

| Cell Line | Clinical Status | Location of muscle | Sex | Age |
| --- | --- | --- | --- | --- |
| **C.1** | Non-MND control | Paravertebral | M | 16 |
| **C.2** | Non-MND control | Paravertebral | F | 13 |
| C.3 | Non-MND control | Quadriceps | M | 38 |
| SMAII.1 | SMA Type II | Paravertebral | M | 11 |
| SMAII.2 | SMA Type II | Paravertebral | F | 7 |
| SMAII.3 | SMA Type II | Lumbar | F | 13 |

We next evaluated the myotube differentiation of each cell line. Myotube formation is a process composed of different steps [29], including cell cycle withdrawal, alignment, fusion, and nuclear organization. Our 2D culture model system allows us to reach the myotube stage with aligned nuclei (**Supplemental Fig.1 F**), in three days (**Figure 1A**). To avoid any confounding effects from the modestly different rates of cycling between the cell lines (**Supplemental Fig.1 G**), the same number of control or SMA myoblasts were seeded to achieve ∼90% confluency, allowed to adhere for several hours, and switched to differentiation media within the day (**Figure 1B**). We observed a decrease in the fusion index, defined as the number of nuclei included in myotubes compared to the total number of nuclei, in SMA myoblasts compared to controls (**Figure 1C**) but no change in the maturation index, the number of myotubes with three or more nuclei (**Figure 1D**). We validated that these changes in fusion were not due to increased cell death in the SMA myoblasts (**Supplemental Fig.1H**).

**Figure 1:**
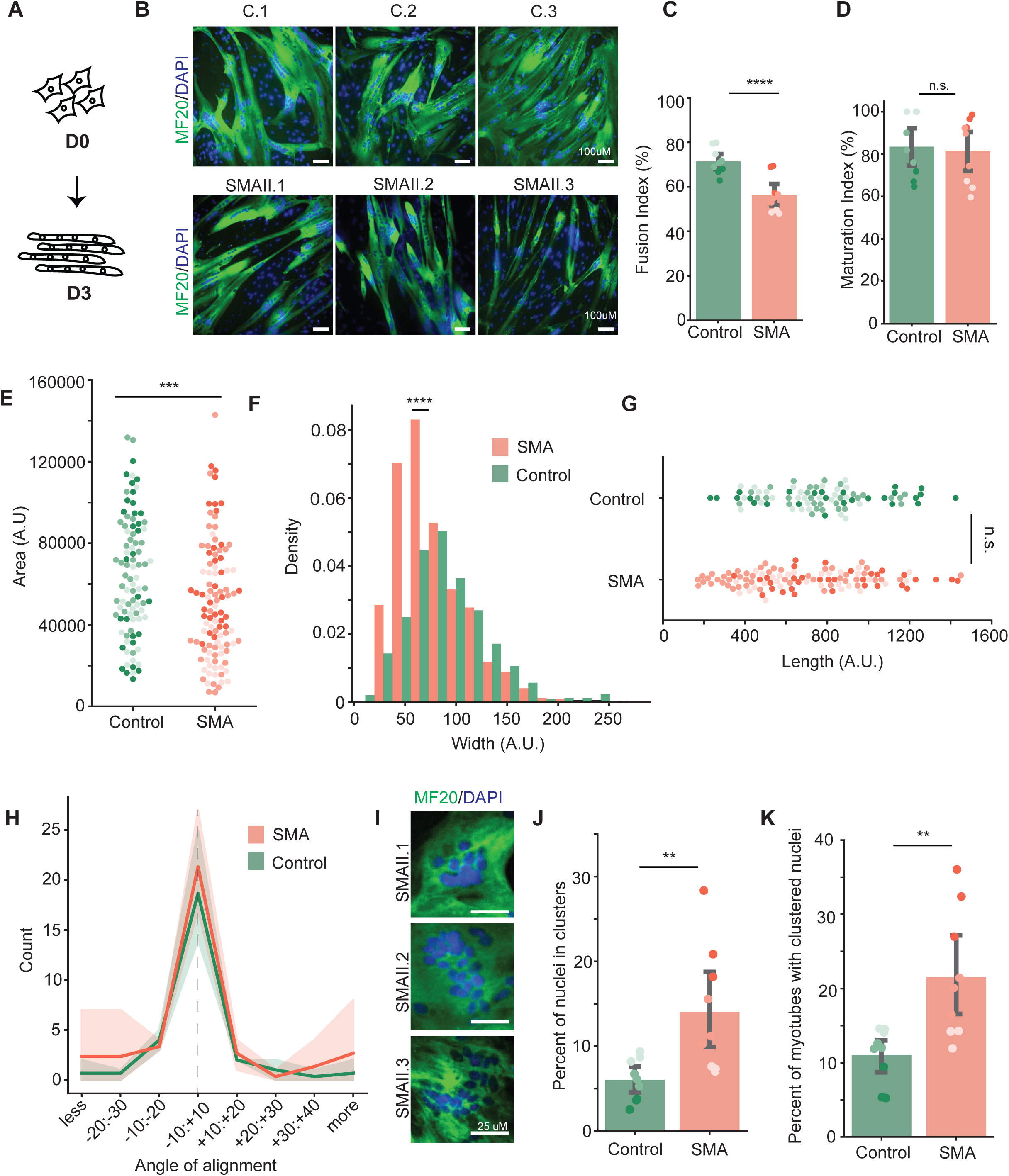
SMA myoblasts show defective and disorganized differentiation. **A)** Schematic of myoblast differentiation in a 2D in vitro culture system. Day 0 refers to the myocyte state when cells are confluent and Day 3 refers to the most mature myotubes that can be generated *in vitro* in controls. **B)** Representative images of the three control and three SMA myotubes stained with MF20 and DAPI. **C)** Quantification of the fusion index in SMA and control myotubes. Each point represents the fusion index for an individual field from a cell line (n=3 replicates for each cell line). Each cell line is colored with a different hue. Means were tested using a student’s t-test. **** p<0.00001. **D)** Quantification of the maturation index in SMA and control myotubes. Each point represents the maturation index for an individual field from a cell line (n=3 replicates for each cell line). Each cell line is colored with a different hue. Means were tested using a student’s t-test. n.s. = not significant. **E)** Quantification of the area of SMA or control myotubes. Each point represents a single quantified myotube, colored by the shade of cell line. Means were tested using a student’s t-test. *** p<0.0001. **F)** Quantification of the distribution of widths in SMA and control myotubes. Means were tested using a student’s t-test. **** p<0.00001. All cell lines are pooled together after individual measurement. **G)** Quantification of the average length of SMA and control myotubes. Each point represents a single quantified myotube, colored by the shade of cell line. Means were tested using a student’s t-test. n.s. = not significant. **H)** Quantification of the alignment of myotubes in control and SMA differentiation. The dashed line represents perfect parallel alignment. Data are binned by how many myotubes show a degree of alignment that is different from parallel alignment. Means were tested using a student’s t-test. n.s. = not significant. All three cell lines are pooled together for this representation. **I)** Representative images of the nuclear aggregation in SMA myotubes. Control myotube nuclear alignment can be found in Supplemental Figure 1F. **J)** Quantification of the percentage of myotubes with aggregated nuclei. Each point represents an individual field from a cell line (n=3 replicates for each cell line). Each cell line is colored with a different shade. Means were tested using a student’s t-test. ** p<0.001. **K)** Quantification of the percentage of nuclei that are found in aggregated clusters. Each point represents an individual field from a cell line (n=3 replicates for each cell line). Each cell line is colored with a different shade. Means were tested using a student’s t-test. ** p<0.001.

To assess the morphology of the myotubes that formed from the SMA and control, we analyzed length, area, alignment, width, and nuclei organization. Defects were found in the total area (**Figure 1E**), width (**Figure 1F**) but not in the length (**Figure 1G**). We also analyzed the alignment of the myotubes and observed no differences (**Figure 1H**). Finally, we analyzed the nuclear organization, which should be aligned along the center of the myotube at this stage [30] and observed that both the percentage of nuclei in clusters (**Figure 1I-J**, **Supplemental Fig. 1F**) as well as the percentage of myotubes with clustered nuclei were increased in SMA compared to control cell lines myotubes compared to controls myotubes with abnormal or clustered nuclei as well (**Figure 1K**). Altogether, this data demonstrated that SMA Type II myoblasts exhibit a reduced fusion into myotubes and a distorted fusion into myotubes.

### Transcriptional and chromatin profiling of SMA myoblasts reveals downregulation of *MYOD1* target genes and increased genomic and stress response pathways

To better understand what molecular mechanisms might lead to an altered myotube fusion profile, we profiled the transcriptome of myotubes after three days of differentiation using RNA-sequencing (**Figure 2A**, **Supplemental Fig. 2A**). Comparing all three SMA myotube cell lines, we observed 180 upregulated and 198 downregulated genes (**Supplemental Fig. 2B, Supplemental Table 1**). Among the downregulated pathways, we had an enrichment for DNA damage response pathways and cell division pathways (**Supplemental Fig. 2C**). Among the upregulated pathways we observed extracellular matrix (ECM) organization, including the upregulation of several collagen proteins (*COL1A1, COL1A2* and *COL6A3*) and *TGFB1* (**Supplemental Fig. 2C, Supplementary Table 1**), which is known to stimulate fibrosis in the context of diseased muscle [31], recalling what we had previously observed in our SMA Type II muscle cohort [10]. Among the most upregulated genes we found *IRF7*, which was originally identified in the context of EBV infection and has emerged as a crucial regulator of type I interferons (**Supplemental Table 1**). We then performed upstream transcription factor (TF) analysis to determine the upstream regulators of the differentially expressed genes in myotubes (**Supplemental Fig. 2D**), which found STAT1, a known effector the interferon response [32] as a putative upstream regulator, as well as a loss of activity of MYOG, which is known to be involved in myocyte fusion [33].

**Figure 2:**
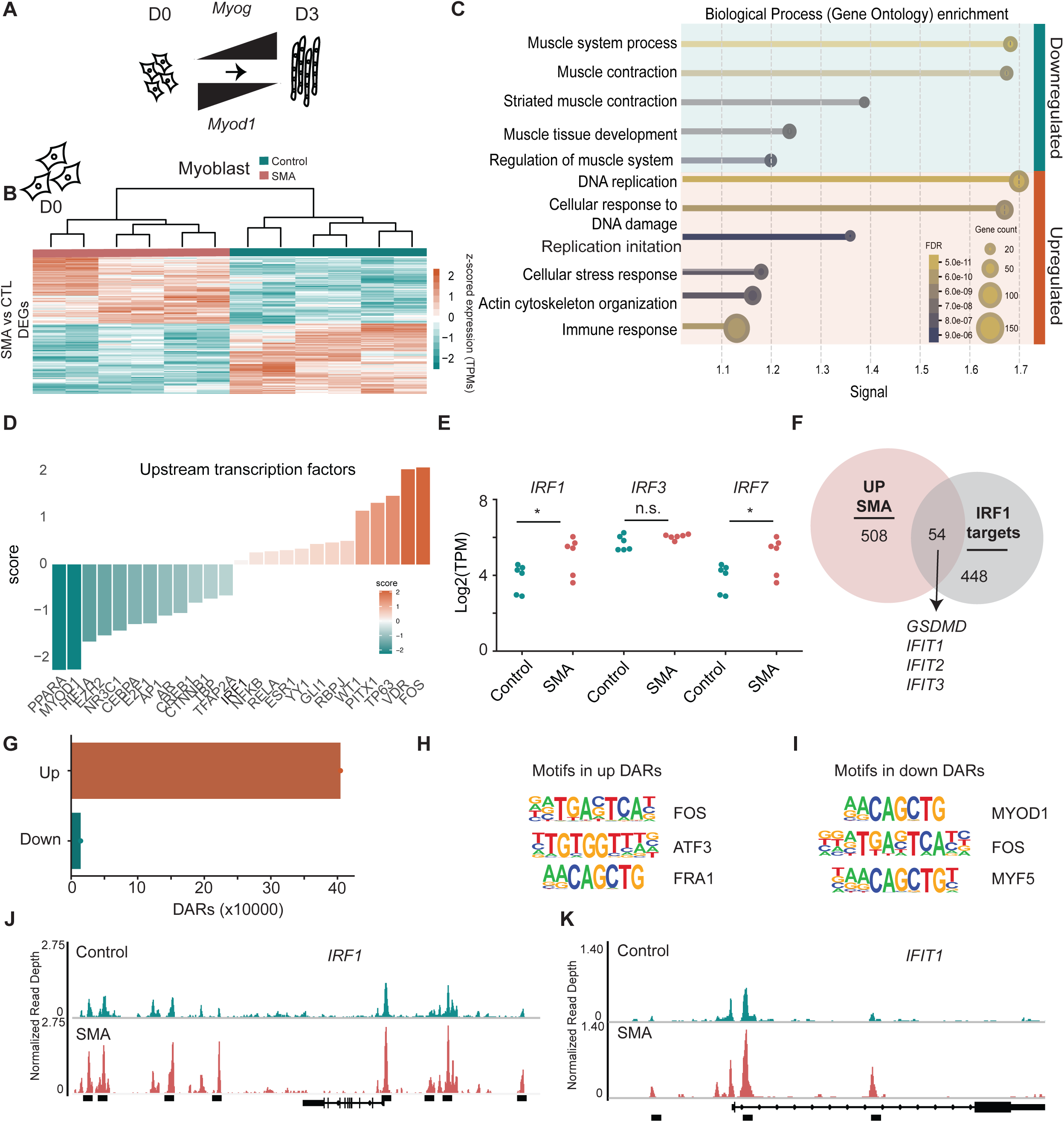
SMA myoblasts have an increased transcriptional and chromatin signature of cellular stress. **A)** Schematic of myogenesis in 2D in vitro culture system, where MYOD1 levels decrease over time as MYOG levels increase over time. **B)** Heatmap of the differentially expressed genes (DEGs) between all SMA myoblasts (n=3 biological replicates) and controls (n=3 biological replicates). **C)** Pathway analysis of the upregulated and downregulated genes from A. **D)** Upstream regulator analysis for the differentially expressed genes between SMA and control myoblasts. Scoring represents the putative activity of the transcription factors (TFs), where positive scores indicate an increase in activity and negative scores indicate a decrease in activity. **E)** Transcript per million (TPM) for IRF1, IRF, and IRF7. Statistics represent the adjusted p-value for DEGs analysis. **F)** Overlap between all IRF1 target genes (taken from the enrichR database) and the upregulated genes in SMA myoblasts. Four example genes from the set of 54 are highlighted. **G)** Bar plot of the number of differentially accessible regions (DARs) between SMA and control myoblasts (combined) as measured by ATAC-seq. **H-I)** Statistically significant motifs in the DARs in the peaks with increased accessibility (H) and decreased accessibility (I). **J-K)** Representative bigwig plots showing reads from control and SMA myoblast ATAC-seq at IRF1 (J) and IFIT1 (K). Significantly different peaks between the two conditions are marked with a black bar.

To better understand if problems leading to the impaired differentiation in the SMA myoblasts started before the fusion, we next characterized the transcriptome of the myoblast cell lines. We performed RNA-sequencing of the myoblasts before differentiation (**Supplemental Fig. 2A**). Differential expression analysis showed 640 downregulated genes and 576 upregulated (**Figure 2B**). Pathway analysis of the downregulated genes showed many processes associated with muscle function, including contraction and differentiation (**Figure 2C**), which are also known to be altered in SMA Type II paravertebral muscle [10]. Additionally, SMN is known to be involved in organizing actin dynamics [34]. Among the downregulated genes in both myoblasts and myotubes, we also observed *Mymx,* a gene involved in the fusion of myocytes into myotubes (**Supplemental Table 1 & 2**) [35]. We performed upstream regulator analysis on the differentially expressed genes and observed that the activity of MYOD1 was predicted to be decreased (**Figure 2D**). Indeed, overlapping a list of MYOD1 target genes, 35% of the downregulated genes in SMA myoblasts were known MYOD1 targets (**Supplemental Fig. 2E, Supplemental Table 3**). Previous reports have suggested that acute loss of SMN causes a decrease in MYOD1 levels as well as its target genes [19]. In our system, neither the RNA levels of *MYOD1* (**Supplemental Fig. 2F**) nor the protein levels (**Supplemental Fig. 2G**) were significantly different between controls and SMA myoblasts. This may suggest differences between human and mouse myoblast cells.

Intriguingly, among the upregulated pathways, we obtained many terms related to cellular stress response (**Figure 2C**). Within these cellular stress response pathways, we could observe two major axes: DNA damage repair processes, including the upregulation of *MRE11*, *BRAC1* and *RAD51*, and innate immune responses, including the upregulation of *CGAS, TMEM173, IRF7, IRF1, IFIT2 and IFIT3, TLR3 and GSDMD* (**Supplemental Fig. 2H, Supplemental Table 2**). This was further supported by upstream regulator analysis, which showed that IRF1 and NFKB targets were activated among the upregulated genes (**Figure 2D**). mRNA levels of both *IRF1* and *IRF7*, but not *IRF3*, were increased in SMA myoblasts (**Figure 2E**) and 10% of the upregulated genes in the SMA myoblasts were predicted to be IRF1 target genes (**Figure 2F**, **Supplemental Table 4**).

To better understand how these transcriptional changes were regulated, we performed ATAC-sequencing to allow us to measure sites of accessible chromatin, often representing enhancers bound by transcription factors [36]. We found 40451 regions with increased chromatin accessibility and 1424 regions with decreased chromatin accessibility (**Figure 2G**, **Supplemental Tables 5-6**). As expected, the majority of either gained or lost peaks were found in distal intergenic regions or introns, two common locations for enhancers (**Supplemental Fig. 2H**). We observed that in the top 15 motifs of peaks that had increased chromatin accessibility, we had FOS, ATF3 and FRA1 (**Figure 2H**, **Supplemental Table 7**), while in the regions with less accessible chromatin in SMA myoblasts, we found an enrichment for the MYOD1 and MYF5 motif (**Figure 2I**, **Supplemental Table 8**). In contrast to the downregulated regions, we found that 5.17% of the peaks contained bZIP:IRF motifs and 3.22% of peaks contained IRF3 motifs (**Supplemental Table 7**). Accessible chromatin was found on IRF1 (**Figure 2J**) and on several IRF1 target genes, including *IFIT1* (**Figure 2K**). Collectively, our transcriptional and chromatin accessibility analysis suggested that cellular stress at the myoblast stage, causing activation of cellular stress pathways and chromatin remodeling, may underlie the differentiation problems in SMA myotubes.

### SMA Type II myoblasts have DNA damage prior to fusion

The accumulation of DNA damage is linked to innate immune activation and the progression of many diseases, including neurodegenerative disorders, neuromuscular diseases, and immune deficiencies[37–39]. Furthermore, in myoblasts, the presence of sustained DNA damage is known to promote the formation of myotubes with a distorted phenotype [40], similar to what we observed in SMA. SMA cell lines had a 2.5-fold increase in the percentage of nuclei with 53BP1 foci, a marker associated with double-stranded break (DSB) resolution (**Figure 3A-B**, **Supplemental Fig.3A**). DNA damage was also confirmed by quantifying the ratio of yH2AX to total H2AX by western blot (**Supplemental Fig.3B**) where we observed a 2-fold increase in the phosphorylated form of the histone H2AX in SMA myoblasts (**Figure 3C**). This data aligns with the p53 activation observed in muscle biopsies from SMA type II patients [10] and the DNA fragmentation detected in the muscle from an SMA mouse model[41,42]. Furthermore, it underlines a cell-intrinsic defect in SMA myoblasts, independently of the denervation caused by motor neuron death. After fusion, SMA myotubes did not show DNA damage (**Supplemental Fig. 3C-D**), consistent with the transcriptomic data showing a downregulation in DNA damage response pathways (**Supplemental Fig.2C**) and in line with previous findings about a decline of DNA damage responses during myogenic differentiation [43].

**Figure 3:**
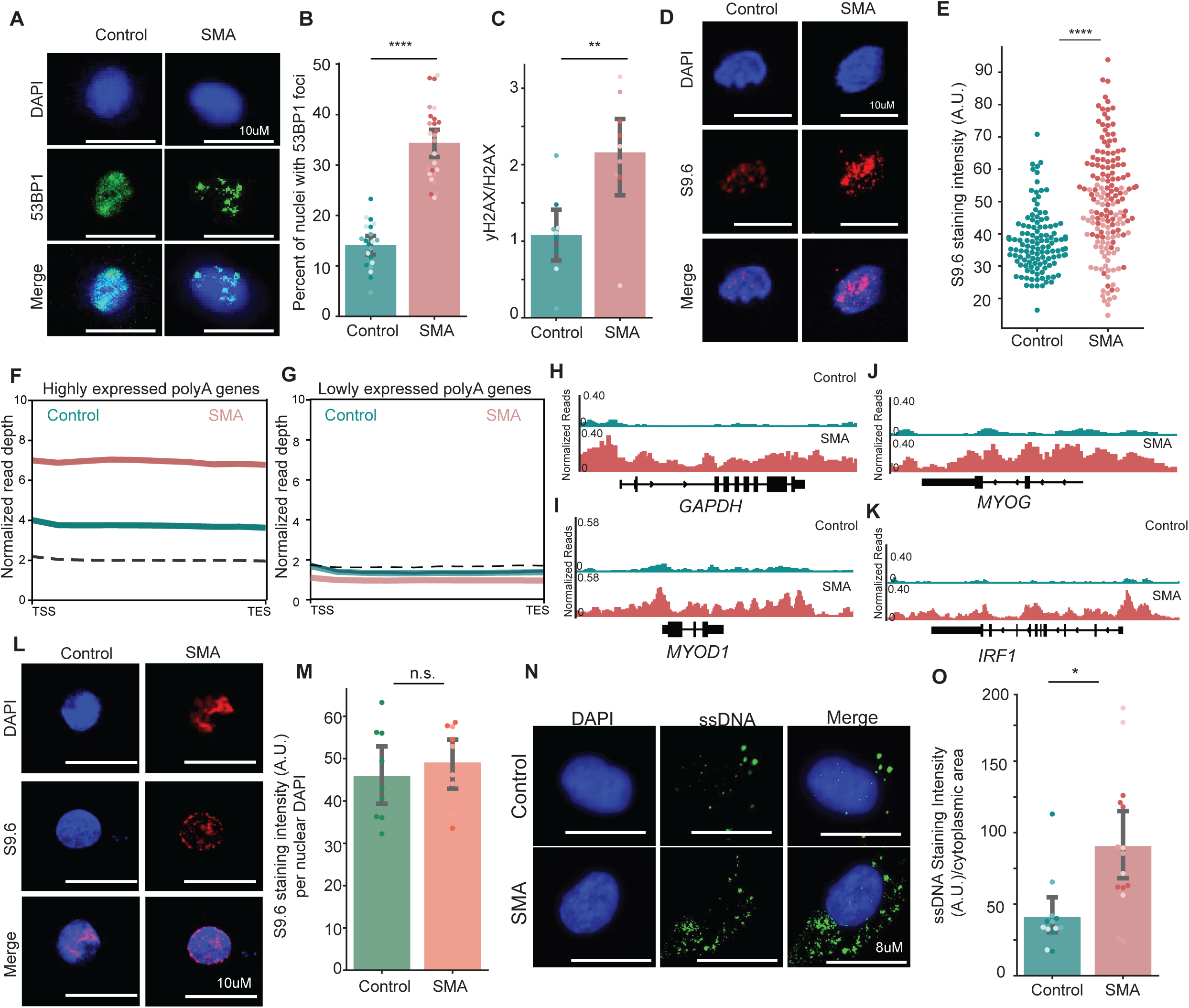
SMA myoblasts have an accumulation of DNA damage and R-loops. **A)** Representative images of 53BP1 staining on one control and one SMA myoblast cell lines. All the cell lines can be found in the supplemental. **B)** Quantification of 53BP1 staining in SMA and control myoblasts. Each cell line is plotted in a different hue. Means were tested using a student’s t-test. **** p<0.00001. **C)** Quantification of Western blot for yH2AX/H2AX. Full western blot is shown in Supplemental Figure 3. Each cell line is plotted in a different hue. Means were tested using a student’s t-test. ** p<0.001. **D)** Representative images of R-loop staining with the S9.6 antibody in nuclei extracted from SMA (SMAII.1) or control myoblasts (C.1). **E)** Quantification of S9.6 staining intensity in control (C.1) vs SMA (SMAII.1 and SMAII.2) nuclei. Each point represents the quantification of a single nucleus in the image. The difference between the means was tested using a student’s t-test. **** p<0.0001 **F-G)** Density plot of reads from S9.6 Cut&TAG mapping the location of R-loops on the highly expressed genes (>1000 TPMs) (F) on the lowly expressed genes (10-50 TPMs) in SMA (SMAII.1 and SMAII.2) and control (C.1 and C.2) cell lines. The dashed line represents the IgG negative control. **H-I)** Representative bigwig plots of the Cut&TAG data from SMA and control myoblasts on housekeeping (H), lineage-specific genes (I-J) and a gene upregulated in only SMA myoblasts (K). **I)** Representative images of S9.6 R-loop staining in nuclei extracted from SMA (SMAII.1) and control (C.1) myotubes. **J)** Quantification of S9.6 staining in SMA (SMAII.1 and SMAII.2) and control (C.1) myotubes. Means were tested using a student’s t-test. Each cell line is represented in a different hue. n.s. not significant. **K)** Representative image of ssDNA staining in SMA and control myoblasts. **L)** Quantification of the ssDNA staining from K. Each cell line is represented in a different hue. Means were tested using a student’s t-test. * p<0.01.

### SMA II myoblasts have R-loop accumulation on highly expressed genes

After validating the presence of genomic stress in SMA myoblasts and its dynamics during the fusion process, we next focused on what might be causing this increase in DNA damage. In SMA motor neurons, it has been proposed that excessive R-loop formation leads to double-strand DNA breaks and the activation of apoptotic pathways, contributing to cell death [26]. R-loops are transient RNA:DNA hybrid structures that form during transcription and generate DNA damage if unresolved [44]. It is known that SMN helps assemble an R-loop resolving complex, including SETX, on the RNA Polymerase II CTD domain, to promote transcription termination and avoid permanent RNAP II stalling. In the absence of SMN, R-loops are not efficiently removed, causing genome instability due to stalled transcriptional complexes crashing into the replication machinery [23,24,27,45].

We proceeded to test for R-loop accumulation in myoblasts by immunostaining with S9.6 antibody, which recognizes DNA:RNA hybrids, despite many known limitations[46–49]. We first performed the staining directly on fixed myoblasts and detected a high cytoplasmic background (**Supplemental Fig.3E**), consistent with the literature [48]. Despite this limitation, we observed increased signal in SMA myoblasts (**Supplemental Fig. 3F**) and we validated that this signal was specific using treatment with RNAseH1, which should digest away the RNA component of the DNA: RNA structure and prevent antibody binding (**Supplemental Fig. 3G-H**). To exclude the possible confounding signal of the cytoplasm, we next extracted the nuclei from control or SMA myoblasts and stained with S9.6 and observed a 1.5-fold increase in S9.6 staining (**Figure 3D-E**). To better localize the genes with increased R-loop deposition, we next employed Cut&TAG sequencing [50] to tag the locations of DNA bound by the S9.6 antibody. We observed that there was an increased accumulation of R-loops on SMA myoblasts in highly expressed RNA Pol II dependent genes (**Figure 3F**) compared to lowly expressed genes (**Figure 3G**). We also found R-loop accumulation on RNA Pol I targets, namely rRNA genes (**Supplemental Fig. 3I**), as previously described in SMA [45]. R-loop accumulation was notable on housekeeping genes such as *GAPDH* (**Figure 3G**), key lineage genes such as *MYOD1* (**Figure 3I**) and *MYOG* (**Figure 3J**), as well as on genes upregulated only in SMA myoblasts, such as *IRF1* (**Figure 3K**).

We had previously observed that DNA damage was resolved in SMA myotubes.

Along similar lines, when we measured S9.6 staining in the nuclei of SMA myotubes, we did not find a significant increase in R-loop accumulation (**Figure 3L-M**). In order to be resolved, R-loops must be cut by a series of RNases and topoisomerases, including *SETX* and *ZPR1*, among others [44]. No increase in their expression was observed in SMA myoblasts or myotubes (**Supplemental Tables 1-2**). Resolution of R-loops can result in the export of RNA: DNA hybrids into the cytoplasm, which can result in the accumulation of both ssRNA and ssDNA. We hypothesized that this increased cytoplasmic export of DNA and RNA species, both from the DNA damage, the R-loop resolution, and likely also from mitochondrial failure [16,19], might lead to an increase in overall cytoplasmic DNA species and thus the downstream innate inflammatory signaling we observed. To confirm this accumulation, we performed immunostaining using an antibody for ssDNA and found a significant increase in SMA myoblasts (**Figure 3 N-O**).

### Activation of cGAS in myoblasts by transfection of double stranded DNA or a chemical agonist phenocopies SMA differentiation alterations

After verifying that cytosolic DNA accumulates in SMA myoblasts, we wanted to test the hypothesis that activation of cytosolic DNA sensors by this DNA was responsible, in part, for the altered SMA myotube fusion and the altered transcriptome. To test the effect of increased cytoplasmic DNA on myoblast differentiation, we transfected control myoblasts with 500bp dsDNA fragments and differentiated them for three days (**Figure 4A-B**). dsDNA treatment resulted in a decrease in the fusion index (**Figure 4C**) and in the maturation index (**Figure 4D**) compared to mock treated myoblasts. Myotube morphology was also altered, with treated myotubes having a decrease in area (**Figure 4E**), width (**Figure 4F**) and length (**Figure 4G**). Treatment with dsDNA also induced the upregulation of *IFNB* mRNA (**Supplemental Fig. 4A-B**) and its secretion (**Figure 4H**). To rule out the possibility that the observed fusion defects were merely a consequence of apoptosis or senescence, we assessed cell death using an LDH cytotoxicity assay (**Supplemental Fig. 4C**) and evaluated senescence through a senescence-associated β-galactosidase assay (**Supplemental Fig. 4D**) and observed no differences between dsDNA and mock treated cells.

**Figure 4:**
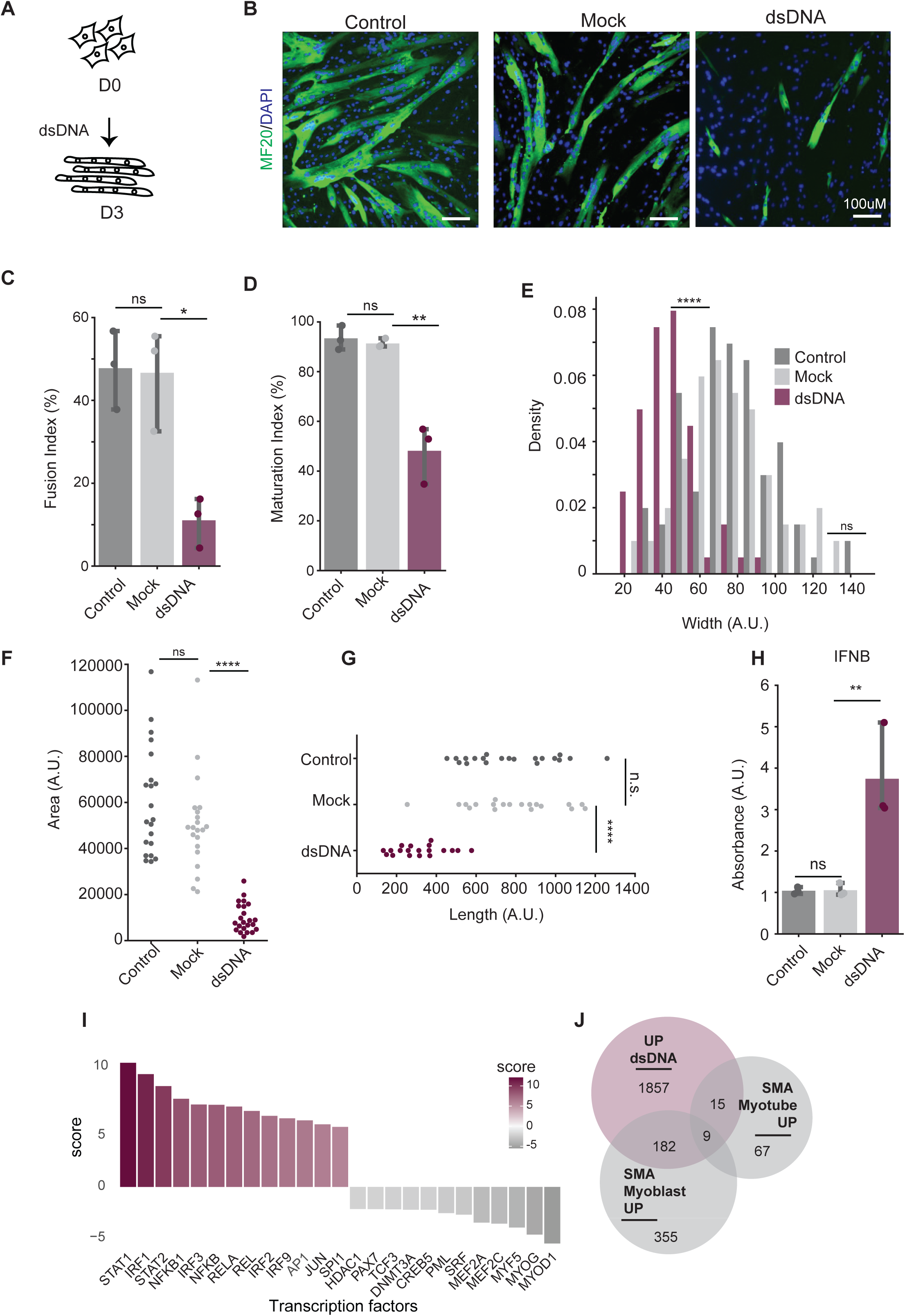
Activation of cytosolic DNA sensing pathways in control myoblasts phenocopies SMA fusion defects and transcriptome. **A)** Scheme of treatment of control myoblasts with dsDNA during the myoblast differentiation process. (n=3 independent treatments for each group) **B)** Representative image of control myoblasts treated with lipofectamine (mock) or dsDNA after 3 days of differentiation. **C)** Quantification of the fusion index of control treated with lipofectamine (mock) or dsDNA after 3 days of differentiation. Means were tested using a one-way ANOVA test. *p<0.01. **D)** Maturation index of control myoblasts treated with lipofectamine (mock) or dsDNA after 3 days of differentiation. Means were tested using a one-way ANOVA test. **p<0.001. **E)** Quantification of dsDNA-treated myotube width. Means were tested using a one-way ANOVA test. ****p<0.00001. **F)** Quantification of dsDNA-treated myotube area. Means were tested using a one-way ANOVA test. ****p<0.00001. **G)** Quantification of dsDNA-treated myotube length. Means were tested using a one-way ANOVA test. ****p<0.00001. **H)** ELISA measurement of IFNB secretion by dsDNA-treated and mock myotubes. Means were tested using a one-way ANOVA test. **p<0.001. **I)** Upstream regulator analysis for the DEGs between dsDNA-treated myotubes and lipofectamine-treated myotubes (mock) after 24 hours of treatment. **J)** Overlap between the genes upregulated in SMA myoblasts and myotubes and the genes upregulated in dsDNA-treated control myoblasts after 24 hours.

To better understand how the transcriptomic changes in dsDNA treated cells compared to those of SMA myoblasts, we performed RNA-sequencing on myoblasts one and three days after a one-time treatment with dsDNA and compared their transcriptome to mocked treated and untreated cells (**Supplemental Fig. 4E**). Most gene expression changes were observed after 24-hours of treatment with 4033 DEGs at D1 (**Supplemental Table 9**). and 593 at D3 (**Supplemental Table 10**). Indeed, many gene expression changes occurring at day 1 returned to near baselines levels at day three (**Supplemental Fig. 4F**). Pathway analysis of the transcriptional changes at day 1 showed an enrichment for processes related to response to viruses, related to the upregulation of the interferon pathway, and a downregulation of muscle development, translation, and actin cytoskeletal organization (**Supplemental Fig. 4G**). Upstream regulator analysis of the gene expression changes at day 1 showed an increase in the activity of IRF1,2, 3 and NFKB (**Figure 4I**)., consistent with the interferon and NFKB response we observed in SMA myoblasts. We also observed a decrease in the activity of MYOD1, MYOG and MYF5 (**Figure 4I**), similar to what we observed in the SMA myoblasts and myotubes. Indeed, dsDNA treatment could recapitulate about 33% of the upregulated genes in SMA myoblasts (**Figure 4J**) and 36% of the downregulated genes (**Supplemental Fig. 4H**), inclidng the downregulation of *MYMX* (**Supplemental Table 9-10**). Finally, we validated the downregulation of *MYOD1* and *MYOG* mRNA on day three after treatment, showing that the presence of dsDNA in the cytoplasm can downregulated the expression of these key myoblast differentiation factors (**Supplemental Fig. 4I**).

We next wanted to determine if the activation of a DNA sensor alone could also phenocopy the observed differentiation defects. Cytoplasmic RNA-DNA hybrids derived from nuclear R-loops accumulation can activate innate immune responses via cGAS-related interferon signaling [51]. cGAS is a well-known DNA sensor, and indeed, many different sources of cytoplasmic DNA accumulation cause cGAS activation [52]. To verify whether increased cGAS activity could impair differentiation in our *in vitro* system, we treated a control cell line (C1.1) with a cGAS-STING agonist, 2’3’-cGAMP, and differentiated them for 3 days (**Figure 5A-B**). Treated myoblasts had a 4-fold decrease in the fusion index (**Figure 5C**), but no change to the maturation index (**Figure 5D**). Treatment also induced morphological alterations of the myotubes, including a decrease in area (**Figure 5E**), width (**Figure 5F**), and length (**Figure 5G**). As expected, cGAS activation turned on the expression of downstream interferon genes, including a 3-fold increase in *IFNB* mRNA production (**Figure. 5H**). Surprisingly, we did not observe increased IFNB protein production, probably due to the short half-life of ∼6 hours [53] of these species, which makes them hard to detect, or as a consequence of interferon beta being only one of many downstream effectors of cGAS signaling (**Figure. 5I**). These morphological changes phenocopied both what we observed with dsDNA treatment as well as in SMA myotubes, suggesting that cGAS activation alone is enough to reproduce parts of the SMA myoblast phenotype.

**Figure 5:**
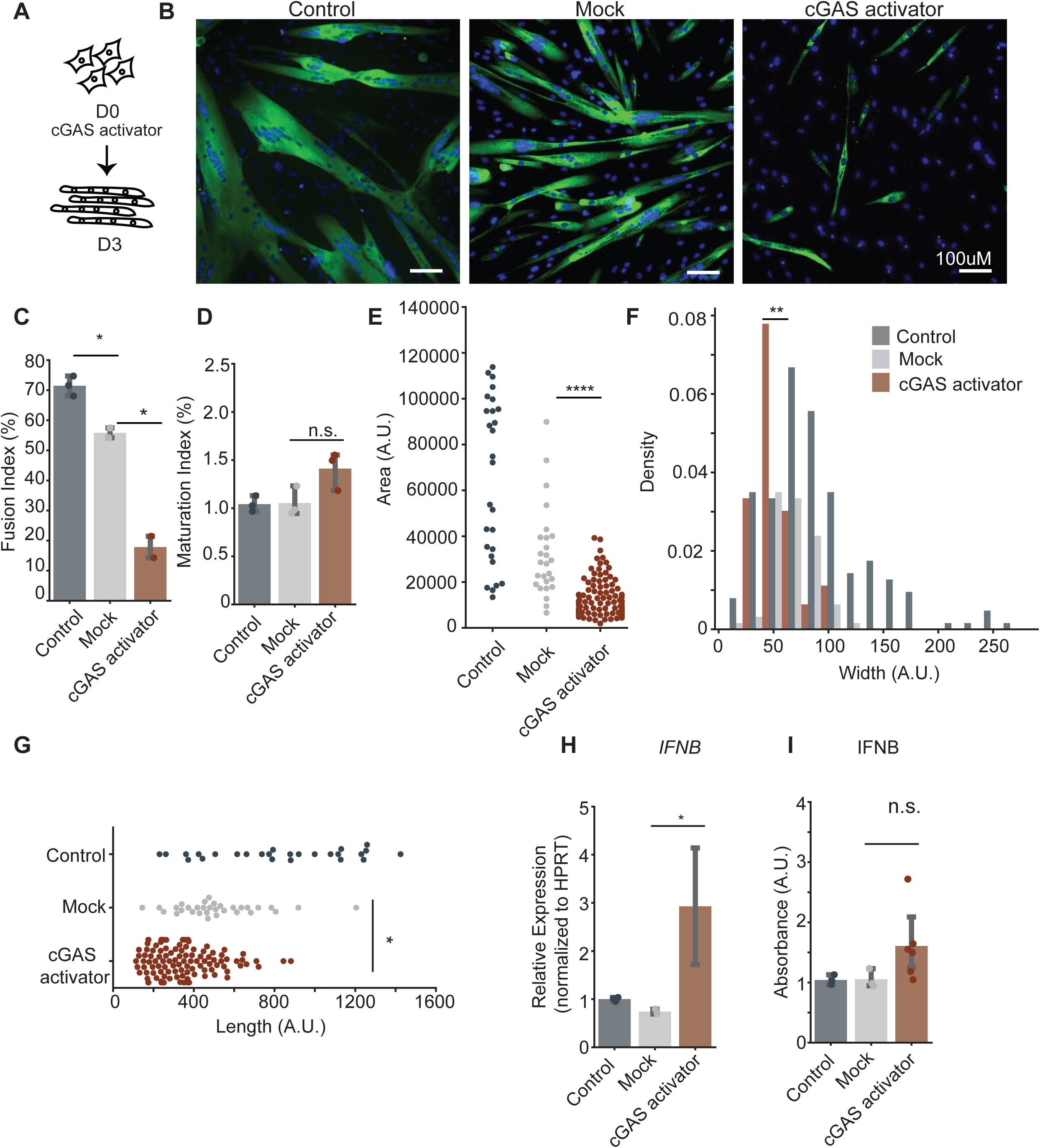
Activation of cGAS DNA sensors causes a fusion defect in control myoblasts. **A)** Scheme of treatment of control myoblasts with cGAS agonist during the myoblast differentiation process (n=3 independent treatments). **B)** Representative image of control myoblasts treated with lipofectamine (mock) or cGAS agonist after 3 days of differentiation. **C)** Quantification of the fusion index of control treated with lipofectamine (mock) or cGAS agonist after 3 days of differentiation. Means were tested using a one-way ANOVA test. *p<0.01. **D)** Maturation index of control myoblasts treated with lipofectamine (mock) or agonist after 3 days of differentiation. Means were tested using a one-way ANOVA test. n.s. not significant. **E)** Quantification of cGAS agonist-treated myotube area. Means were tested using a one-way ANOVA test. ****p<0.00001. **F)** Quantification of cGAS agonist-treated myotube width. Means were tested using a one-way ANOVA test. **p<0.001. **G)** Quantification of cGAS agonist-treated myotube length. Means were tested using a one-way ANOVA test. *p<0.01. **H)** qPCR for IFNB expression in cGAS agonist-treated myotubes. Means were tested using a one-way ANOVA test. *p<0.01. **I)** ELISA measurement of IFNB secretion by cGAS agonist-treated and mock myotubes. Means were tested using a one-way ANOVA test. n.s. not significant.

### Treatment with H-151 STING inhibitor improves SMA myotube differentiation

Cytosolic DNA sensors have several downstream effectors, the majority of which are mediated by the endoplasmic reticulum (ER) membrane protein STING (*TMEM173*) [54]. To test the hypothesis that increased activation of cytosolic DNA sensors in SMA myoblasts was the cause of impaired myoblast fusion, we treated SMA myoblasts with a STING antagonist (H-151) every 24 hours in differentiation media for 3 days (**Fig. 6A-B**). Treatment increased the fusion index to that of control myoblasts (**Fig. 6C**) and increased the average width of SMA myotubes (**Fig. 6D**) but did not alter the area or length (**Supplemental Fig. 5A-B**). We did not observe any changes in the mRNA or protein levels of IFNB after H-151 treatment (**Fig. 6E**, **Supplemental Fig. 5C**). Collectively, these data support that dampening the downstream signaling from STING can improve SMA myotube fusion. However, the permanence of unresolved differentiation defects suggests that there may be other cytosolic DNA sensors and, therefore, innate immune pathways involved.

**Figure 6:**
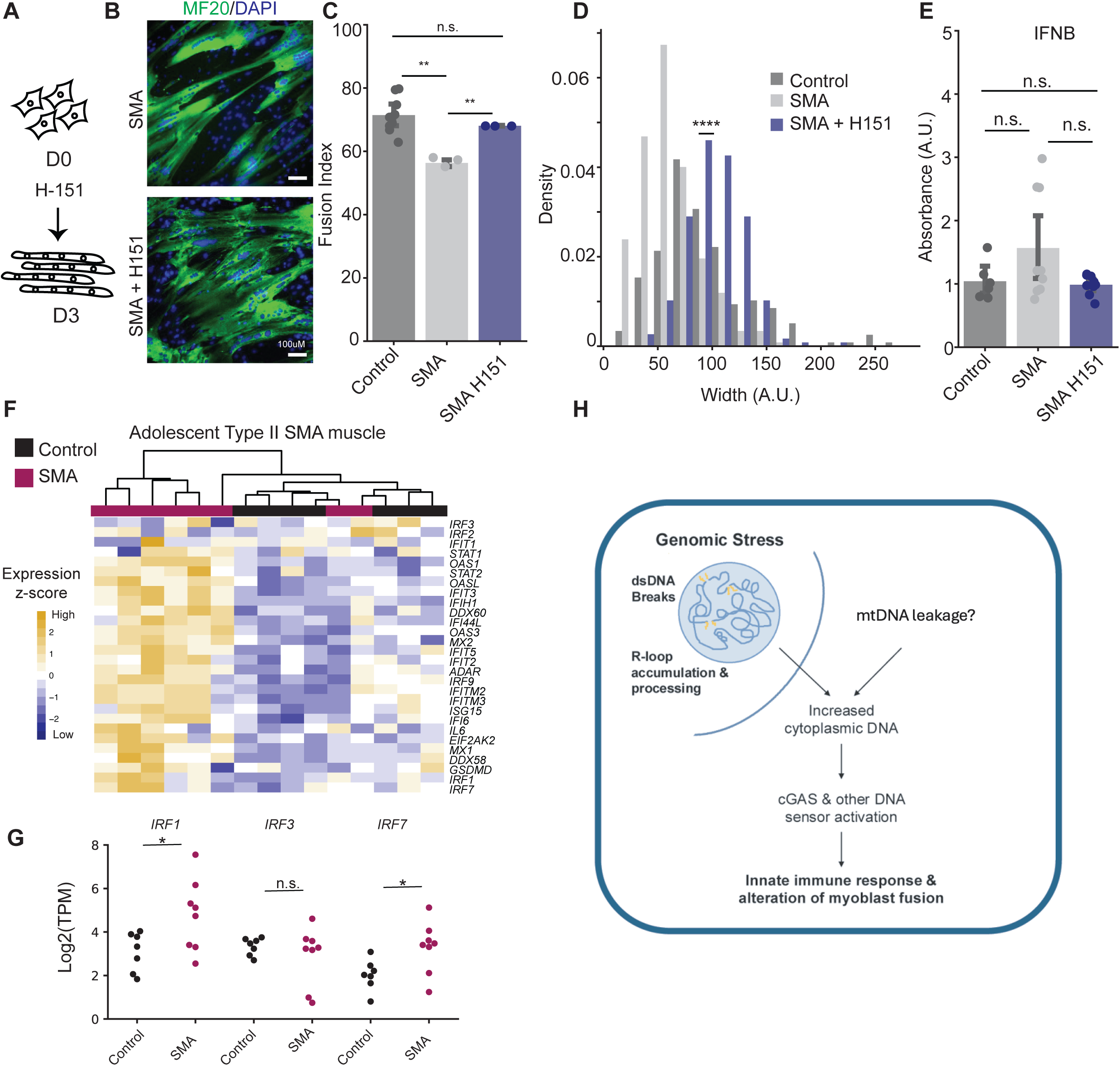
Inhibition of STING improves SMA myoblast fusion. **A)** Scheme of treatment of SMA myoblasts, cell line SMAII.1, the most severely affected, with the STING inhibitor H-151 during the myoblast differentiation process (n=3 independent treatments per group). **B)** Representative image of SMA myoblasts treated with H-151 inhibitor. **C)** Quantification of fusion index in H-151-treated SMAII.1 myoblasts. Means were tested using a one-way ANOVA test. **p<0.001. **D)** Quantification of the width of H-151-treated myoblasts (SMAII.1) and control (C.1). Means were tested using a one-way ANOVA test. ****p<0.00001. **E)** ELISA measurement of IFNB secretion by H-151-treated myotubes. Means were tested using a one-way ANOVA test. n.s. not significant. **F)** Heatmap of interferon pathway and innate immune signaling in paravertebral muscle from Type II SMA patients and controls. Data reanalysed from Grandi et al., 2024. **G)** TPMs of *IRF1*, *IRF3*, and *IRF7* from that data in F. **H)** Schematic of the proposed summary model for the role of cytosolic DNA sensing in SMA, causing alterations in myoblast fusion and differentiation.

### Interferon pathways are upregulated in SMA Type I and Type II muscle samples

Our *in vitro* results suggested that loss of *SMN1*, and a reduction in total SMN protein levels in myoblasts engender an activated innate immune response, including the activation of cytosolic DNA sensing pathways, which affect the myoblast fusion and differentiation process. Indeed, fusion defects have already been observed in Type I SMA fetal muscle [11], and Type I interferon signaling was observed in Type I mouse SMA models [55]. To assess whether the pathways identified in our SMA Type II myoblasts were also activated in muscle from SMA Type II patients, we used our previously published RNA-sequencing data [10] and analyzed the same target gene list, observing that 6 of the 8 (75%) Type II patients had an activation of the STING pathway (**Fig. 6F**). Notably, our previously histological analysis of these same muscles had not found increased immune cell infiltration, suggesting that this signature might come from the muscle tissue itself. Further, as in the myoblasts, we observed an increase in *IRF1* and *IRF7* mRNA levels, but not of *IRF3* (**Fig. 6G**). This was further validated in a separate RNA-sequencing dataset [56] of fetal or perinatal Type I diaphragm and iliopsoas muscle (**Supplemental Fig. 5D-E**). Altogether, both datasets in combination with our *in vitro* study suggest that genomic stress due to *SMN1* loss in SMA causes an increase in cytoplasmic DNA in myoblasts, leading to increased innate immune activation and consequential differentiation defects (**Fig. 6H**).

## DISCUSSION

Spinal Muscular Atrophy (SMA) is a complex, multifactorial disease [2]. The early and primary symptoms are caused by the death of motor neurons, causing paralysis and eventual death without treatment. However, increasing evidence from clinical reports and animal studies highlights the involvement of other tissues, including skeletal muscle.

However, a clear understanding of the precise role of *SMN1* in specific tissues remains unclear. In this work, we performed an *in vitro* study of the effects of *SMN1* loss on myoblast cells, the early progenitors of muscle tissue. We characterized the ability of Type II SMA myoblasts to differentiate into myotubes and described a role for SMN loss-induced genomic stress in activating innate immune signaling in SMA myoblasts. We show that this activation of innate immune signaling, be it through dsDNA or direct activation of cGAS, contributes to the disorganized and defective differentiation of myoblasts. Altogether, this work highlights a new consequence of SMN loss in muscle cells and adds to our understanding of the complexity of SMA pathology.

### Disorganized myoblast differentiation is a feature of SMA in vitro

SMA Type II myoblasts exhibited impaired differentiation dynamics, forming smaller myotubes with less width and total area than controls. SMA myotubes exhibited reduced fusion and increased nuclear clustering. These data suggested either a alteration in the differentiation, with SMA myotubes appearing immature compared to the controls, or a defect in the differentiation process, where SMA myotubes failed to complete differentiation properly and form abnormal myotubes. In agreement with these data, previous studies on satellite cells and myoblasts derived from skeletal muscles of SMA mouse models have demonstrated comparable myogenic impairments, including reduced myotube size and altered expression of differentiation markers [57,58]. Similarly, *Smn* knockdown (*Smn*-KD) in the C2C12 cell line led to the formation of malformed myotubes, which were smaller and often mononucleated, demonstrating that the myogenesis defects are directly correlated to SMN protein deficiency [18]. To date, few studies have been conducted on human cells *in vitro*, and the majority have focused on the most severe viable form of SMA, Type I. Myoblasts derived from human muscle biopsies (SMA Type I patients) or SMA-hESC lines (SMA Type I-like) confirmed the myogenic impairment, showing reduced myoblast fusion and lower formation of multinucleated myotubes [16,17]. Remarkably, a similar result was obtained *in utero* from histological analysis of embryonic muscle biopsies from SMA type I fetuses at 12 weeks of gestation, showing smaller, thinner, and not equally distributed myotubes, suggesting an intrinsic role of SMN loss in muscle development [55]. Moreover, smaller myofibers have also been found in muscle biopsies of patients affected by infantile spinal muscular atrophy [59].

It is important to consider that SMA is a highly heterogeneous disease, as evidenced by the variable symptom severity experienced by patients and variability in treatment responses[10]. Indeed, within our patient derived cell lines, we could see a range of fusion impairments, suggesting other cellular pathways may act as modifiers. Additionally, we cannot rule out that the immortalization process may influence some of the pathways under investigation, such as those involved in apoptosis and DNA damage repair. However, taken together with the literature, our data add to the evidence that SMA myogenic progenitors, at least in Type I and Type II patients, have reduced differentiation capacity and validate a cell-intrinsic role of SMN in myoblasts and myotube biology. Indeed, our myoblast transcriptome analysis also demonstrated the upregulation of genes involved in cellular stress response, DNA damage, DNA replication, and immune response. We hypothesized that the transcriptomic changes observed at the myoblast stage drive the disorganized and defective myotube formation.

### Loss of SMN in myoblasts causes genomic stress

Our work also reinforces the conclusion that one of the major effects of SMN loss in the muscle cells is an increase in genomic stress. An increase in DNA damage has already been observed in several SMA tissues and *in vitro* models, including fibroblasts, motor neurons, muscle, and spinal cord [26]. In SMA motor neurons, the accumulation of double-strand breaks (DSBs) throughout the genome is proposed to trigger cell death pathways, either through p53 activation or JNK-mediated degeneration, contributing to their death [27,60,61]. Embryonic muscle biopsies from SMA type I fetuses at 12 weeks of gestation showed evidence of DNA fragmentation in myotubes, but in contrast to motor neurons[62–64], no signs of apoptosis were observed [11]. Similarly, in our study, SMA type II myoblasts exhibited DNA damage in the absence of cell death or increased markers of senescence. Put together, these observations highlight the complexity of SMA phenotypes, and how the loss of the same protein can lead to similar molecular consequences - namely the accumulation of DNA damage – but with different cellular outcomes due to their intrinsic machinery.

Indeed, muscle cells are known to be efficient at repairing DNA damage without always leading to apoptosis and cell death [42,65,66,43]. This difference in cell response may help to explain the differences in how SMA pathology presents in the central nervous system versus peripheral tissues.

The presence of DNA damage in SMA is often correlated with R-loop accumulation [26]. R-loops are three-stranded structures formed by a DNA:RNA hybrid and a displaced single-stranded DNA (ssDNA) [67]. When improperly regulated, as is the case in SMA [26], they interfere with DNA replication, repair, and transcription, causing genome instability, including DNA damage [68]. Accumulation of R-loops has been reported in several diseases, including other neurodegenerative diseases [69]. Pathological R-loop accumulation in SMA has been observed in fibroblasts from SMA Type I patients, in primary spinal cord neurons from SMA mice (*SMN*Δ*7)*, in SMA type I patient spinal cord and mouse tissues (*SMN*Δ*7*), and in SMA iPSC-derived motor neurons [45,70]. Now we confirm that their accumulation is also found in myoblasts and is correlated with increased DNA damage, both of which are resolved after myotube formation.

### Sterile inflammation caused by genomic stress is part of SMA muscle pathology

Genomic stress, including DNA damage, has increasingly emerged as a cause of sterile inflammation, triggered by the presence of DNA or RNA species in the cytoplasm of the cell [39]. For example, R-loops are known immunogenic species; their resolution on the genome triggers the release of DNA/RNA fragments into the cytoplasm, leading to chronic inflammation [51,71–73]. These fragments, whether as ssDNA, DNA: RNA hybrids, or dsDNA, can be recognized by cGAS and other cytosolic DNA sensors, thereby triggering the inflammatory cascade and activating innate immune responses. We found increased ssDNA fragments in the cytoplasm of SMA Type II myoblasts compared to controls, and the transcriptomic analysis revealed the upregulation of innate immune pathways associated with *IRF1* and *IRF7* expression. Similarly, transcriptomic analysis of muscle biopsies from SMA type I and type II patients showed increased innate inflammation. Such innate immune signaling was previously proposed by a mouse study of Type I SMA muscle [55], but a detailed description and characterization were not undertaken. As such, this work adds a new dimension to SMA muscle biology, showing that innate immune activation and interferon signaling are part of the SMA myoblast phenotype.

Increased interferon signaling has been described to play a role in other myopathies with muscle atrophy, including idiopathic inflammatory myositis and myotonic dystrophy type I (DM1) [74–77], particularly as the aberrant interferon I production impairs myoblast differentiation and induces atrophy of myotubes *in vitro* [78]. Therefore, we hypothesize that chronic innate immune activation in SMA myoblasts could explain part of the differentiation defects observed in SMA myotubes (**Figure 6H**). Treatment with dsDNA, more stable than DNA: RNA hybrids or ssDNA, in a control myoblast cell line recapitulated the differentiation impairment, suggesting that part of the SMA phenotype is explained by inflammation.

Previously, Mankan et al. discovered that cytosolic DNA: RNA hybrids activate the cGAS-STING pathway, causing inflammation [79]. Treatment of the same control myoblast cell line with a cGAS agonist also mimicked the SMA differentiation defects, confirming that innate immune inflammation plays a role in SMA muscle pathology. Moreover, as a proof of concept, we treated an SMA cell line with a STING inhibitor, H-151. We observed improvements in certain differentiation parameters, such as myotube fusion and width, while others, like area, remained unaffected. This supports the involvement of the STING pathway in the pathology but also suggests that additional innate immune pathways may contribute to the disease mechanism. The exact list of DNA sensors and interferons involved in SMA pathogenesis remains to be studied. Moreover, the cytoplasmic DNA may have multiple origins, including DNA damage from various sources, the resolution of R-loops, as well as from the leakage of mitochondrial DNA, as mitochondria a known to be damaged in SMA muscle[10,13].

Collectively, the findings of this study provide evidence for cell-intrinsic roles of SMN in muscle, suggesting that the muscle defects observed in SMA patients are not only a consequence of denervation and motoneuron loss. This, together with our previous work on SMA muscle biopsies [10]. which showed transcriptomic remodeling, mitochondrial defects, and DNA stress pathways after Nusinersen treatment, stresses the need for combinational therapies targeting the peripheral organs. Moreover, it offers new insights about the role of SMN in SMA muscle pathology, demonstrating that SMN loss induces genomic stress and subsequent activation of the innate immune signaling.

## MATERIALS AND METHODS

### Myoblast isolation, culture, and differentiation

Immortalized myoblasts from controls (AB1190, male, 16 years old, paravertebral muscle, KM1421, female, 13 years old, paravertebral muscle, and AB1079, male, 38 years old, quadriceps) and SMA (KM432, male, 11 years old, paravertebral muscle, KM1150, female, 7 years old, paravertebral muscle, and AB1141, female, 13 years old, lumbar muscle) muscle were obtained from the MyoLine Platform (Myology Institute, Paris, France). Cells were cultured at 37°C and 5% CO2 under humidity in myoblast proliferation medium (1 volume Medium 199 [Thermo Fisher Scientific, 41150020] + 4 volumes DMEM [Thermo Fisher Scientific, 61965-026] + 20% FBS [Gibco]) supplemented with fetuin (25 μg/mL; Life Technologies, 10344026), hEGF (5 ng/mL; Life Technologies, PHG0311), bFGF (0.5 ng/mL; Life Technologies, PHG0026), insulin (5 μg/mL, Sigma-Aldrich, 91077C-1G), and dexamethasone (0.2 μg/mL, Sigma-Aldrich, D4902). Cells were passaged with 0.05% trypsin (Thermo Fisher Scientific) and routinely tested for mycoplasma (Lonza). For any assays performed in the myoblast stage, cells were taken in the proliferation phase (∼50%–60% confluent). For differentiation assays (myotubes), cells were plated at 90% confluence on plates previously coated with Matrigel (Corning) diluted at 1:20 in media at 37°C for 1h, and allowed to attach for 2–5 hours, before changing to differentiation media (DMEM + 10 μg/mL insulin). Myotubes were collected after 3 days of differentiation.

### Determination of SMN copy number

Genomic DNA was extracted from approximately 200.000 myoblasts with the DNeasy Blood and Tissue Kit (Qiagen). The relative copy number of SMN1 and SMN2 was determined using a previously established set of primers [80]. Briefly, the primers were designed to distinguish between SMN1 and SMN2 in exon 7 at position 6 and intron 7 at position +214. SMN1: forward 5′-TTTATTTTCCTTACAGGGTTTC-3′ and reverse 5′-GTGAAAGTATGTTTCTTCCACGTA-3′. SMN2: forward 5′-TTTATTTTCCTTACAGGGTTTTA-3′ and reverse 5′-GTGAAAGTATGTTTCTTCCACGCA-3′.

For the relative normalizing gene, RPPH1 (forward 5′-CGCGCGAGGTCAGACT-3′ and reverse 5′-GGTACCTCACCTCAGCCATT-3′) was used, also previously established. DNA was diluted to 10 ng/μL in water, and 2 μL was loaded into a 25 μL SYBR Green reaction with 0.05 μM primer. Cycling conditions were modified from the original to 50°C for 2 minutes, 95°C for 10 minutes, 45 cycles of 95°C for 15 seconds, and 58°C for 1 minute. Samples were normalized using the RRHP1 housekeeping gene, and the relative copy number was determined using the ΔΔCt method. The absolute copy number of SMN2 in the SMA cell lines was measured using digital droplet PCR (Bio-Rad) according to the manufacturer’s instructions. TaqMan probe for the combined SMN1/2 was purchase from ThermoScientific (Hs04979908_g1 FAM) and albumin was used as a loading control (F-GCTGTCATCTCTTGTGGGCTGT, R- ACTCATGGGAGCTGCTGGTTC, P-CCTGTCATGCCCACACAAATCTCTCC) using the VIC fluorophore. 50ng of DNA were loaded in each ddPCR reaction using the ddPCR Supermix for Probes (BioRad) according to the manufacturer’s instructions. The copy number was calculated using the ddPCR software on double positive droplets.

### RNA isolation and library preparation for RNA-sequencing

RNA from cells was collected using the RLT buffer from the RNeasy Plus Mini Kit (Qiagen) according to manufacturer’s instructions and frozen at -80C until ready for extraction. RNA extraction was performed according to the manufacturer’s instructions. RNA quantity was established using a Nanodrop spectrophotometer and RNA integrity number (RIN) was calculating using a BioAnalyzer (Agilent Tapestation 4200). All samples had RIN score greater than 9. mRNA library preparation was performed using poly A enrichment and sequenced on the Illumina NovaSeq sequencer using paired-end 150bp reads.

### RNA-sequencing analysis and differential gene expression

The sequencing libraries were processed using the nf-core RNA pipeline[81] (https://nf-co.re/rnaseq/usage) using the standard parameters. Reads were mapped to the mouse genome (mm10). The resulting gene counts were determined using Salmon[82] and used for downstream analysis with DeSeq2 [83] Metascape [84], and enrichR [85]. Differential expression analysis was done using DeSeq2, using the following parameters: abs(log2FoldChange) > 0.5 & padj <=0.01 & lfcSE <0.5). Upstream factor analysis for the transcription factors that controlled a given gene set was performed using the decoupleR, which allows transcription factor activity to be inferenced from bulk RNA-sequencing data using a list of known motifs [86]. These transcription factors are given a score based on how active or inactive they are in the list of differentially expressed genes. All RNA-sequencing data was visualized using custom R-scripts.

### Quantitative Real Time PCR

1 μg of RNA extracted from different cell lines was converted into cDNA using the High-Capacity cDNA Reverse Transcription Kit (Applied Biosystems) according to the manufacturer’s recommendation. PCR reactions (20 μL) were set up in duplicate using TaqMan Mix (Applied Biosystems) with SMN1/2 probe (Hs00165806_m1), Myogenin probe (Hs00231167), MyoD1 probe (Hs00159463), IFNβ probe (Hs01077958), and HPRT (Hs02800695_m1) as a housekeeping control gene. Relative expression was calculated using the ΔΔCt method.

### Assay for Transposase-accessible chromatin (ATAC)- sequencing and analysis

ATAC-seq libraries were prepared from myoblast cell lines as previously described[36]. Libraries were sequenced on a NovaX sequencer (Illumina) at a depth of 10 million read pairs per libraries. Libraries were prepared from two cell lines for each genotype (C.1, C.3 and SMAII.1, SMAII.2). Peaks were called using MACS2[87] and peak sets were merged per genotype and then between genotypes using the previously described peak merging algorithm[36]. From the combined peak set, differentially accessible peaks were called using DeSeq2[83]. Motif analysis was performed using HOMER[88]. Peaks were annotated to their closest gene using ChipSeeker [89]. Peaks were visualized as bigwig files using IGV [90].

### Cleavage Under Targets and Tagmentation (CUT&Tag) sequencing and analysis

CUT&Tag was performed using the iDeal CUT&Tag Kit for Histones for chromatin profiling kit (Diagenode) with a few changes from the manufacturer’s instructions. 300’000 myoblasts per CUT&Tag reaction were used. Before bead binding, cells were fixed in 0,1% methanol-free PFA at room temperature for 5 minutes, and the reaction was quenched using 50 μL of Glycine 1.25M for 5 minutes. 12 μg of S9.6 (Merck #MABE1095) antibody per reaction was used. The quality of the library was assessed by Agilent Tapestation, while the quantity was determined by NEB Quantification kit for Illumina (#E7630SL). Libraries were sequenced on a NovaX Illumina sequencer to a read depth of 10 million read pairs per library. Reads were mapped to the hg38 genome using HiSat2[91] and duplicate reads were removed using deeptools[92]. Bam files were then converted to bigwig files using a BamCoverage from deepTools and used to perform enrichment analysis across different genomic regions using computeMatrix and plotHeatmap from deepTools [92]. Highly and lowly expressed genes were determined using the RNA-sequencing data from above. A gene was considered highly expressed if it was above 500 transcripts per million (TPM) in all myoblasts samples and lowly expressed if it was between 10-50 TPM in each myoblast sample.

### Protein extraction and Western blotting

500,000 myoblasts or myotubes were lysed in RIPA buffer (Sigma-Aldrich) supplemented with a protease inhibitor cocktail (Complete Mini, Roche Diagnostics) and a phosphatase inhibitor (Sigma-Aldrich). Proteins were quantified using the DC Protein Assay (BioRad), and 40 μg of protein was loaded. Proteins were heated to 100°C for 5 minutes and run in 4%–12% Bis-Tris gels (Bio-Rad) and transferred to membranes using the Turbo Transfer System (Bio-Rad). The membranes were then incubated with the corresponding antibody: anti-SMN (1:1,000; BD Biosciences, 610647), anti-γH2AX Ser139 (1:1,000; Cell Signaling #9718), anti-H2AX (1:1,000; Cell Signaling #2595), or anti-vinculin (1:1,000; Sigma-Aldrich, V9191), diluted in Tris-buffered saline containing 0.2% Tween 20 (TBST, Bio-Rad) supplemented with 5% non-fat dry milk. Primary antibody incubation was performed overnight. After 3 washes in TBST, the membranes were incubated with a horseradish peroxidase-conjugated secondary antibody (1:10,000; GE Healthcare) diluted in the blocking buffer. The membranes were further developed using the chemiluminescence substrate SuperSignal Ultra reagent (Thermo Fisher Scientific) or the SuperSignal West Femto Maximum Sensitivity Substrate (Thermo Fisher Scientific) and imaged on a Bio-Rad ChemiDoc MP imaging system. The signal was quantified using ImageLab.

### Edu Labelling

Cell division in SMA versus control myoblasts was assayed using the Click-iT Plus Edu Alexa Fluor 488 Flow Cytometry Assay Kit (Invitrogen) according to the manufacturer’s instructions. Briefly, myoblasts were seeded at 30-40% confluence and subsequently treated with the provided Edu solution at a concentration of 20uM for 6 hours. Cells were then labelled with the Click-IT kit as specified and fluorescence was measured using the BD CytoFlex Flow Cytometer (Beckman) and the percentage of labelled cells was calculated for each cell line.

### Immunofluorescence

Cells were fixed in 4% paraformaldehyde for 15 min at room temperature and washed 3 times in PBS 1X. Myotubes were permeabilized with 0.3% Triton X-100 (BioRad) in PBS 1X, blocked in blocking solution (2% bovine serum albumin (BSA), 1% normal goat serum (NGS) in 0.3% Triton X-100 in PBS 1X) for 1 hour at room temperature and incubated with primary antibody overnight at 4°C (1:100 MYH1E Antibody #MF 20 Hybridoma bank). Cells were incubated with Alexa Fluor–conjugated secondary antibodies (Invitrogen). Nucleus was counterstained with Hoechst 33342 (ThermoFisher) for 10 minutes. Myoblasts were permeabilized with 0.1% Triton X-100 (BioRad) in PBS 1X, blocked in blocking solution (2% bovine serum albumin (BSA) in 0.1% Triton X-100 in PBS 1X) for 1 hour at room temperature, and incubated with primary antibody overnight at 4°C (1:100 anti-53BP1 Cell signalling #49375, 1:75 S9.6 Merck #MABE1095, 1:50 anti-DNA Antibody, single stranded specific, clone F7-26 Merck #MAB3299). Cells were incubated for 1 hour at room temperature with Alexa Fluor–conjugated secondary antibodies (Invitrogen). Nucleus was counterstained with Hoechst 33342 (ThermoFisher) for 10 minutes. To obtain the RNaseH1 control for the R-Loop staining, before fixation, myoblasts were treated with 2U of RNaseH (NEB #M0297S) in PBS and RNaseH buffer for 20 minutes at room temperature. The staining was visualized on the BioTek Lionheart FX Automated Microscope. Fluorescence images were collected with a camera on the BioTek Microscope System or Olympus FV1200 Confocal Laser Scanning Microscope.

### Myotubes differentiation assessment and fusion/maturation index measurements

Human myoblasts were fully differentiated after 3-day induction, if kept in culture for longer, the myotubes detach from the culture dish. ImageJ (88) was used to quantify the area, the length, the width, and the alignment of MF20+ cells in arbitrary units. Images were all acquired on the BioTek Microscope System or Olympus FV1200 Confocal Laser Scanning Microscope. Individual whole myotubes were traced manually and quantified using the “measure” function. Fusion index was measured using the software MyotubeAnalyzer [93] as the percentage of the number of nuclei in MF20+ cells relative to the total number.

Maturation index was measured by manual counting as a percentage of myotubes with two or more nuclei (considered as “mature”) versus the total number of myotubes. All analyses were performed in a blinded manner.

### Double stranded DNA (dsDNA) treatment

Myoblasts were transfected at 80% confluence with dsDNA of indicated lengths using Lipofectamine 2000 (Invitrogen) in differentiation media according to the manufacturer’s instructions. Mock transfection indicates lipofectamine treatment without DNA. Double-stranded DNA molecules were generated by PCR amplification with DreamTaq polymerase (Thermo Scientific) using pcDNA3.1 as a template. One forward primer was used for all PCRs, and varying reverse primers resulted in different length amplicons. The PCR products were purified using the NucleoSpin Gel and PCR Clean-up, Mini kit (Macherey-Nagel). The size specificity of the PCR products was verified by 4% TAE gel electrophoresis. DNA of different lengths was pulled 1:1 to create DNA stocks (50ng/ul), and 10ul was used for transfection. The following primers were used for the generation of dsDNA molecules from the pcDNA3.1 vector template: Forward primer: 50-CGATGTACGGGCCAGATATACG-30 Reverse primer 300 bp: 50-TCAATAGGGGGCGTACTTGGCA-30 Reverse primer 500 bp: 50-AACTCCCATTGACGTCAATGGGG-30 Reverse primer 836 bp: 50-GCAACTAGAAGGCACAGTCG-30.

### Treatment with cGAS-activator or H-151

Myoblasts were transfected at 80% confluence with 1mg/mL cGAS activator (2’3’-cGAMP Invivogen #tlrl-nacga23-02) using Lipofectamine 2000 (Invitrogen) in differentiation media according to the manufacturer’s instructions. Mock transfection indicates lipofectamine treatment without DNA. Myoblasts were treated with 10 uM STING antagonist H-151 (Invivogen #inh-h151) at 80% confluence in differentiation media, the media was changed, and the inhibitor was re-added fresh every 24 hours for 3 days.

### Senescence and cell death assays

To assess senescence, we used the Mammalian beta-Galactosidase Assay Kit (ThermoFisher) following the “procedure for Adherent Cells in a Microplate” instructions. To assess Cell death, we used the CyQUANT^TM^ LDH Cytotoxicity Assay-Fluorescence Kit (ThermoFisher #C20300) following the manufacturer’s instructions.

### IFN**β** ELISA

Myoblasts were cultured for 2 days at 40% confluence in 200 μL of growth media or for 3 days at 80% confluence in 500 μL of differentiation media. Media was collected and spun at 4°C for 10 minutes at 13’000 rpm. 50 μL of media was used for each reaction. We performed the ELISA (Bio-Techne SAS #DIFNB0) accordingly to the manufacturer’s instructions.

### Statistical analysis and plotting

All plots were generated using python and the seaborn package[94] using custom scripts. Statistics were calculated using python’s scipy.stats package[95] and custom scripts. All means between two groups were compared using a student’s t-test and means between several groups were calculated using a one-way ANOVA, as noted in each legend. Figures were generated using Adobe Illustrator.

### Data availability

Raw fastq files are available on GEO. Differentially expressed genes are provided in Supplemental Table 1.

### Author contributions

S.M. - designed the research study, performed experiments, analyzed, and interpreted data, generated figures, and wrote the manuscript

F.G. – designed the research study, performed experiments, analyzed, and interpreted data, generated figures, and wrote the manuscript.

A.B. - derived the immortalized cells used in this study.

P.S. – designed the research study, oversaw data acquisition and interpretation, and wrote the manuscript.

## ACKNOWLEDGEMENTS

We thank the patients and their families for consenting to the donation of tissue samples to this research, without which the generation of SMA and control myoblast cell lines would not be possible. We would also like to thank the members of the Smeriglio lab for their helpful discussion. We would like to the platforms of MyoBank and Myoline (Institut de Myologie, Paris, France), which make the collection and generation of patient-derived myoblast cell lines possible. In particular, we would like to thank Alexis Filachet who helped in the derivation of SMA cell line SMAII.3. The authors would like to acknowledge the MyoImage Platform of the Center of Research in Myology (Institut de Myologie, Paris, France) which supports the microscopy center and the maintenance and utilization of the microscopes used to generate all the images. We would like to thank the staff of the iGenSeq Platform (Institute du Cerveau, Paris, France) for their help with the quality control of the sequencing libraries and the use of their facilities and Agilent TapeStation device. We would also like to acknowledge the Institut Francaise de Bioinformatic (IF) which provides the HPC server used to perform all bioinformatic analysis in this work.

---

**Supplemental Figure 1: SMA myoblasts show defective and disorganized differentiation.**

**A)** qPCR on genomic DNA extracted from control and SMA myoblasts to determine the SMN1 copy number. Means were tested using a student’s t-test. * p<0.01. **B)** qPCR on genomic DNA extracted from control and SMA myoblasts to determine the SMN2 copy number. Means were tested using a student’s t-test. ** p<0.001. **C)** Digital droplet determined total SMN copies measured on genomic DNA extracted from control and SMA myoblasts. Means were tested using a student’s t-test. * p<0.01. **D)** Representative western blot for SMN protein levels. **E)** Quantification of SMN protein levels in myoblasts compared to vinculin loading reference. Each cell line is represented in a different shade within the main bar graph. Means were tested using a student’s t-test. ** p<0.001. **F)** Representative images of nuclear alignment in control myotubes. **G)** Quantification of the percentage of Edu+ labelled cells after 6 hours of incubation in control and SMA myoblasts. Each cell line is represented in a different shade within the main bar graph. Means were tested using a student’s t-test. * p<0.01 **H)** Quantification of absorbance of ELISA for LDH assay. Each cell line is represented in a different shade within the main bar graph. A lysed cell positive control as well as an LDH standard are included for scale. Means were tested using a student’s t-test. n.s. = not significant.

**Supplemental Figure 2: Transcriptional characterization of SMA myotubes and myoblasts.**

**A)** Principal component analysis (PCA) plot of RNA-sequencing data from control and SMA myoblasts and myotubes. **B)** Heatmap of the differentially expressed genes (DEGs) between all SMA myotubes (n=3 biological replicates) and controls (n=3 biological replicates). **C)** Pathway analysis of the DEGs in SMA myotubes versus controls. **D)** Upstream regulator analysis for the differentially expressed genes between SMA and control myotubes. Scoring represents the putative activity of the transcription factors (TFs), where positive scores indicate an increase in activity and negative scores indicate a decrease in activity. **E)** Overlap between putative MYOD1 targets (from EnrichR) and the downregulated genes in SMA myoblasts. **G)** Western blot (top) and quantification (bottom) of MYOD1 protein levels in SMA myoblasts versus controls. Means were tested with a student’s t-test. n.s = not significant. **H)** STRING network analysis of the genes associated with cell stress responses. Pathway analysis was then performed on this list, and annotations are presented in red, blue, green, and yellow colours. Lines represent the strength of the known genetic interactions between two genes. All depicted genes are differentially expressed between SMA and control myoblasts. **I)** Annotation of peaks with increased or decreased chromatin accessibility in SMA versus control myoblasts.

**Supplemental Figure 3: S9.6 marked R-loops increase in SMA myoblasts.**

**A)** Representative images of 53BP1 staining across all control and SMA myoblast cell lines.

**B)** Representative images of yH2AX/H2AX Western blot in control and SMA myoblast cell lines. **C)** Representative images of yH2AX/H2AX Western blot in control and SMA myotube cell lines. **D)** Quantification of western blots for yH2AX/H2AX in SMA and control myotubes. Means were tested using a student’s t-test. n.s. not significant. **E)** Staining with the S9.6 antibody for R-loops on whole myoblasts. **F)** Quantification of R-loop staining in nuclei in whole staining. Means were tested using a student’s t-test. * p<0.01. **G-H)** Representative images and quantification (H) of staining with S9.6 antibody on SMAII.1 myoblasts treated or not with RNAseH. Means were tested using a student’s t-test. * *p<0.001. **I)** Density plot of reads from S9.6 Cut&TAG mapping the location of R-loops on ribosomal genes in SMA and control cell lines. The dashed line represents the IgG negative control.

**Supplemental Figure 4: Treatment of control myoblasts with dsDNA turns on interferon signaling.**

**A-B)** qPCR for IFNB mRNA after treatment with dsDNA relative to HPRT housekeeping genes after 24hours (A) or 72 hours (B). **C)** ELISA of absorbance for LDH assay in myoblasts after treatment with dsDNA. Means were tested using a one-way ANOVA test. n.s. not significant. **D)** ELISA of absorbance to measure β-galactosidase activity in myoblasts treated with dsDNA. Means were tested using a one-way ANOVA test. n.s. not significant. **E)** Principal component analysis (PCA) plot of RNA-sequencing data from control myoblasts treated with dsDNA for 24 or 72 hours. Each point represents one sequenced sample. (n=2 independent treatments for each group). **F)** Heatmap of all differentially expressed genes at Day 1 (24hours) and Day 3 (72 hours) after treatment with dsDNA. Each row represents a single gene. **G)** Pathway analysis of the DEGs in myoblasts treated with dsDNA after 24 hours. **H)** Overlap between the genes downregulated in SMA myoblasts and myotubes and the genes downregulated in dsDNA-treated control myoblasts after 24 hours. **I)** qPCR for *MYOD1* and *MYOG* mRNA levels after treatment with dsDNA relative to *HPRT* housekeeping genes after 72 hours. Means were tested using a one-way ANOVA test. **p<0.001.

**Supplemental Figure 5: Modulation of cytosolic DNA sensors in SMA myoblasts.**

**A)** Quantification of area in SMA myoblasts treated with H-151. Means were tested using a student’s t-test. n.s. not significant. **B)** Quantification of the length of SMA STING inhibitor-treated myoblasts. Means were tested using a student’s t-test. n.s. not significant. **C)** qPCR for *IFNB* mRNA after treatment with STING inhibitor. Means were tested using a one-way ANOVA test. n.s. not significant. **D)** Heatmap of interferon pathway and innate immune signaling in fetal and perinatal muscle from Type I SMA patients and controls. Data reanalyzed from Auslander et al, 2020. **E)** TPMs of *IRF1*, *IRF3* and *IRF7* from that data in D.

