## Supplemental Figures for "Genomic stress drives activation of interferon signaling and innate immune pathways during SMA Type II myoblasts differentiation"

MAZZUCCHI et al. – Supplemental Figures

Supplemental Figure 1

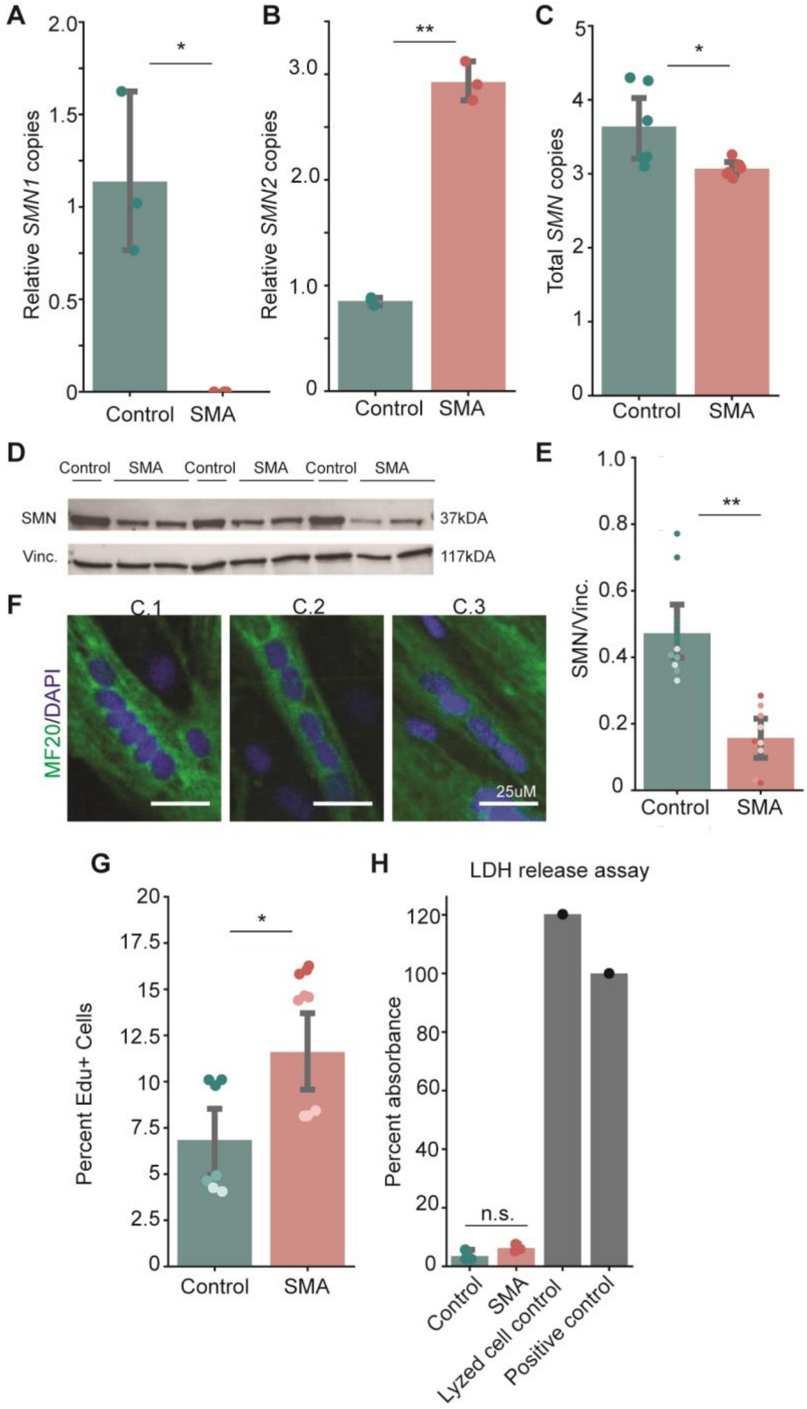

Supplemental Figure 2

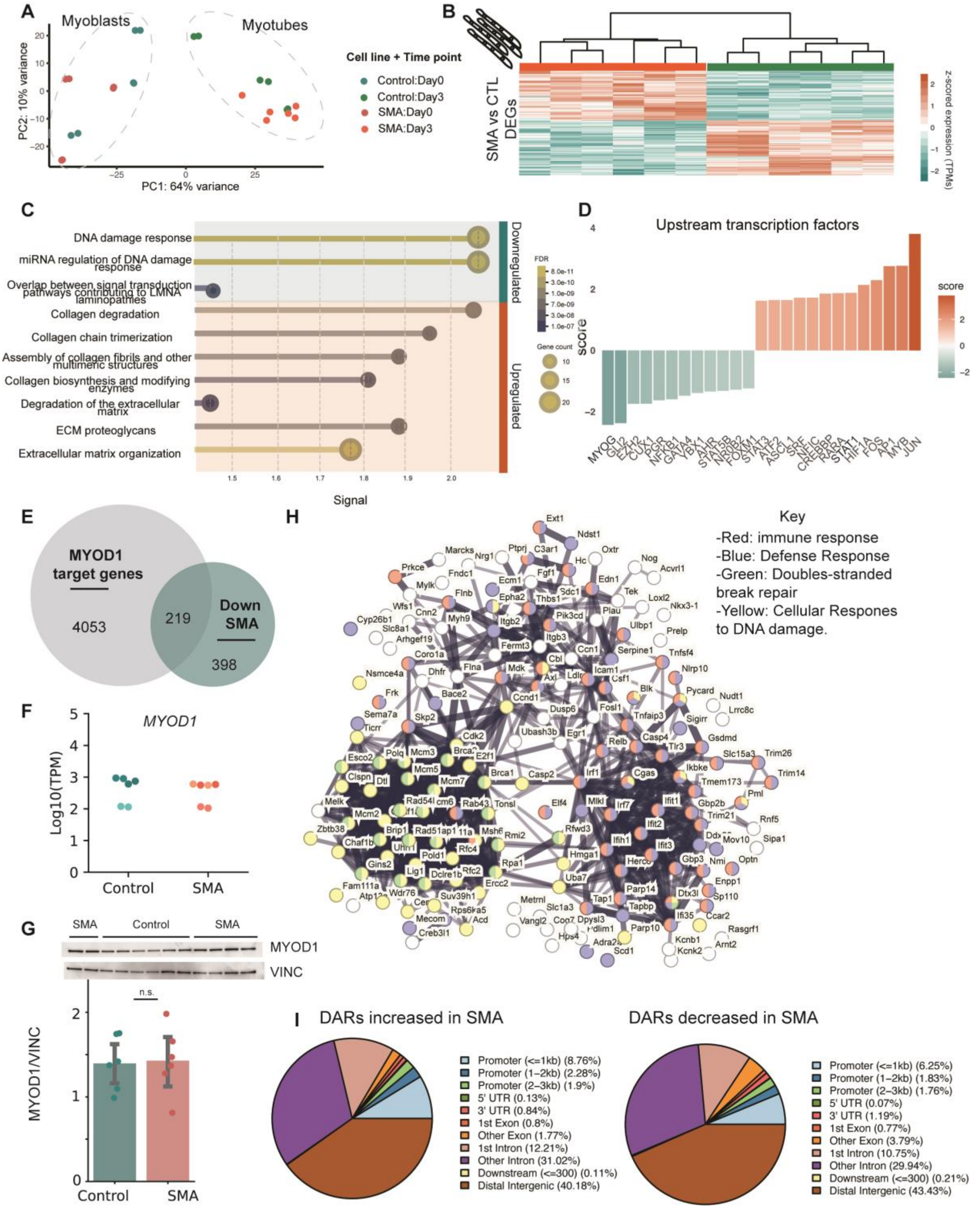

Supplemental Figure 3

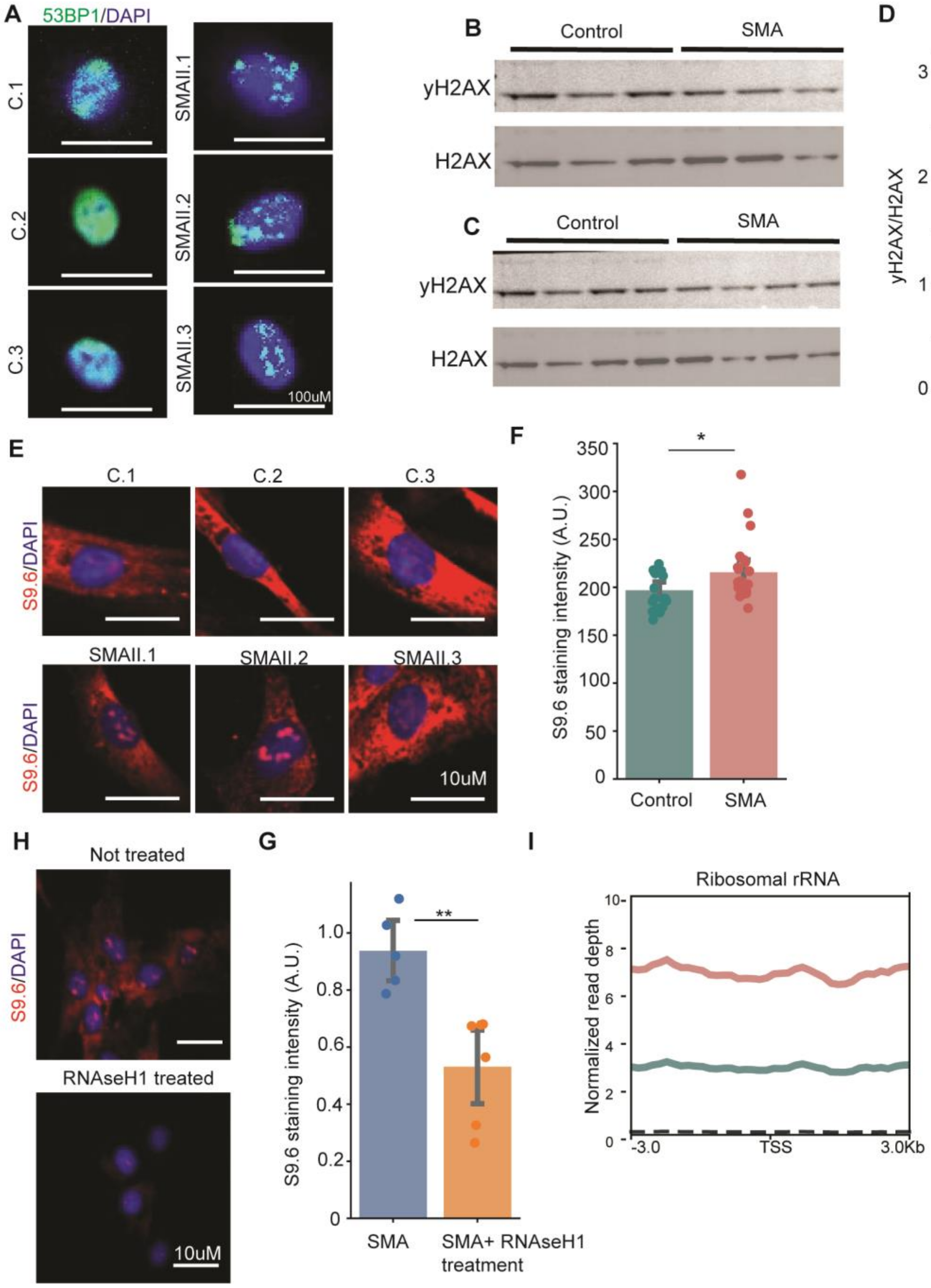

Supplemental Figure 4

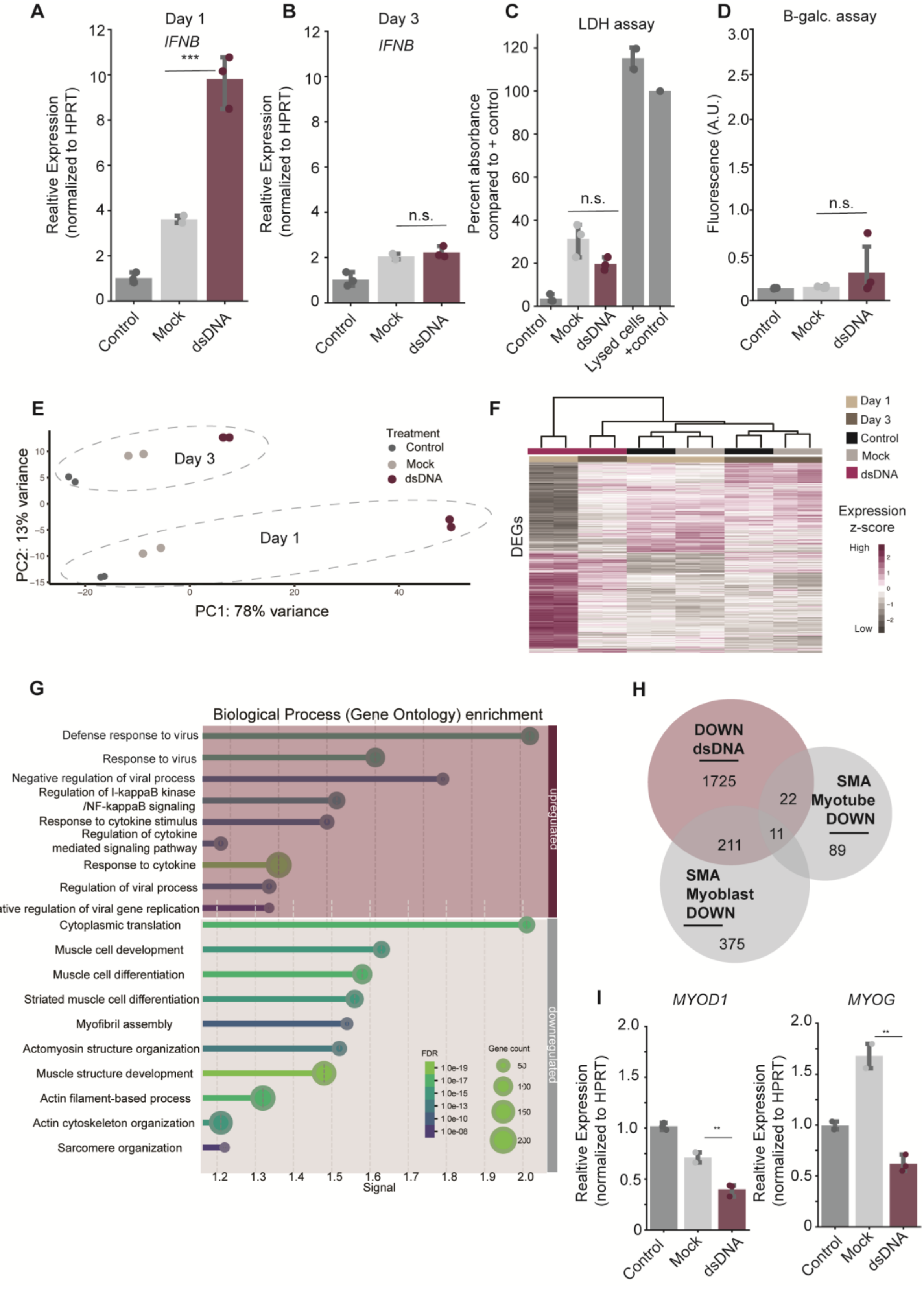

Supplemental Figure 5

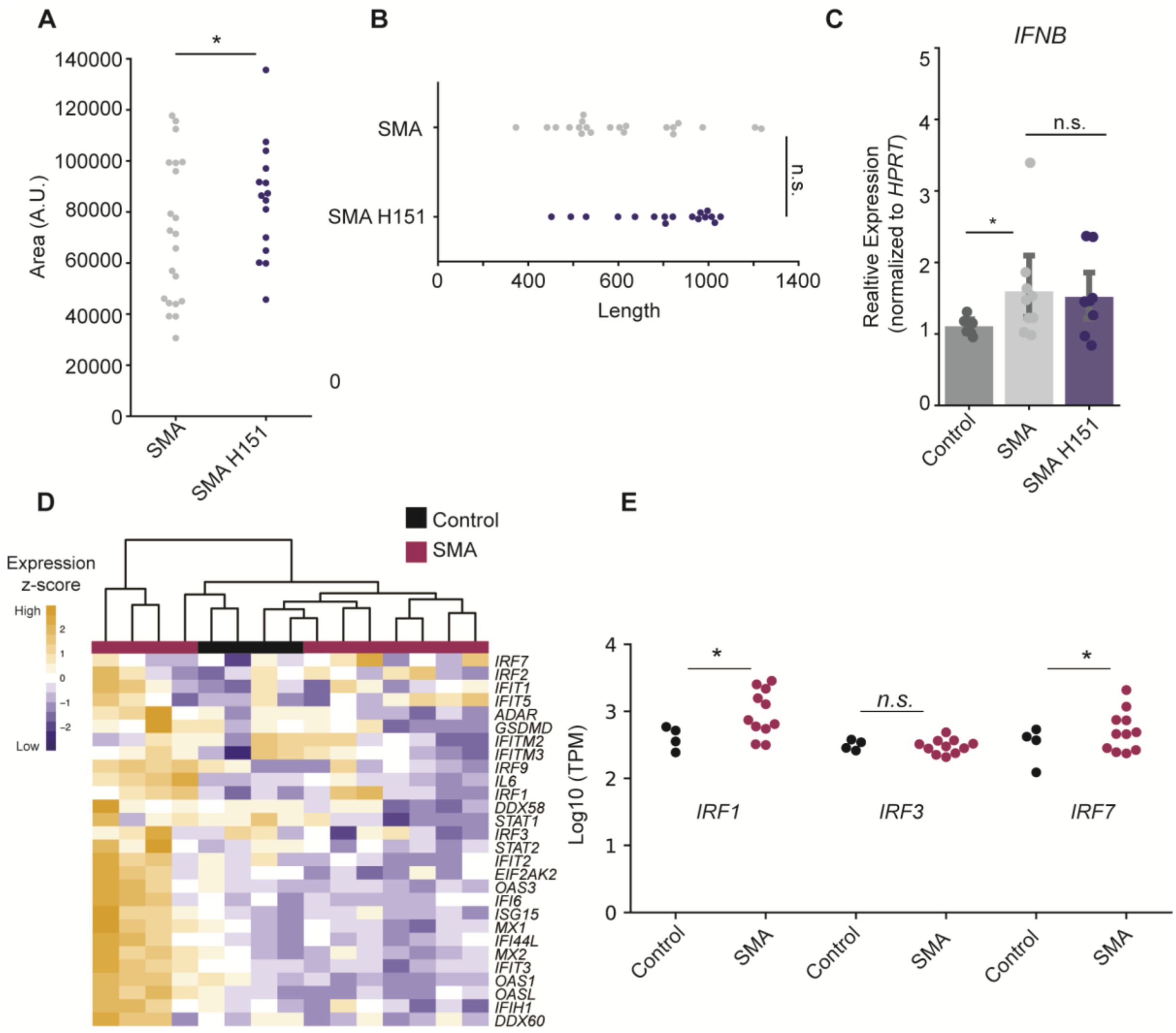
